# Extreme cooling enables survival in extreme heat

**DOI:** 10.64898/2026.07.22.740203

**Authors:** Joanna M. Feehan, Mark C. Bitter, David B. Lowry, Thomas D. Sharkey, Seung Y. Rhee

## Abstract

Complex, multicellular extremophiles face intense challenges in coordinating their cells, tissues, and organs to function in harsh environments. *Tidestromia oblongifolia* is a desert plant that thrives in Death Valley, where air temperatures exceed 49 °C. Using natural collections of *T. oblongifolia* seed, we identified individuals that survive in daily air temperature regimens at the eukaryotic upper thermal limit of 60 °C. Genome-wide association testing detected genetic variation for survival in extreme heat, and transcriptomics implicated pathways regulating leaf cooling. High-throughput infrared imaging revealed that survival was enabled by leaf cooling far below air temperature, identifying a physiological mechanism of thermophily in a multicellular organism. These discoveries provide insights into mechanisms of extreme heat adaptation that could be leveraged to engineer heat-resilient crops.

## Main Text

Most thermophiles are bacteria or archaea with upper thermal limits more than 100 °C (*1–3*). However, thermophilic eukaryotes also exist, with upper thermal limits of ∼60 °C (*4–7*). Thermophilic eukaryotes are often single-celled or filamentous algae and fungi (*8*). Yet, there are unique examples of more complex, multicellular thermophiles such as Pompeii worms living in deep-sea hydrothermal vents (*9*), or panic grass in symbiosis with a virus and fungus growing near geyser basins in Yellowstone (*10*). Knowledge of the molecular mechanisms for thermal adaptation in eukaryotes generally includes thermostability of membranes, protein folding and amino acid substitutions (*8*). It remains unclear if complex thermophilic eukaryotes complement molecular mechanisms with tissue- and organ-level adaptations as part of an adaptive strategy to extreme heat.

Regulation of leaf temperature (*T*_leaf_) may represent such a physiological strategy in plants. *T*_leaf_ is set by the leaf’s energy balance through absorbed radiation, sensible heat exchange with the surrounding air, and latent heat loss through transpiration (*11*, *12*). *T*_leaf_ can therefore decouple from air temperature (*T*_air_), becoming warmer or cooler than *T*_air_ depending on that balance (*13–16*). This is most pronounced in plant species native to extreme biomes (*17*). At high *T*_air_, transpirational cooling can hold *T*_leaf_ below lethal limits, where a difference of only a few degrees can determine survival (*18*).

*Tidestromia oblongifolia* is a plant native to Death Valley, California (*19*) where *T*_air_ set the all-time record of 56.7 °C in 1913 (*20*) and regularly exceeds 49 °C (*21*). This herbaceous perennial thrives in high heat, reaching its maximum growth rate at 44 °C and doubling its biomass roughly every three days (*22*). It depends on access to moisture in subsurface soil layers, but does not store water (*19*, *23*, *24*). Its photosynthetic rates are acclimated to high heat, with optimal carbon assimilation at 47 °C (*19*, *25*). However, as thermotolerance and photosynthesis uncouple at extreme temperatures (*26–29*), we examined heat tolerance independent of photosynthesis. Whether *T. oblongifolia T*_leaf_ matches *T*_air_ in extreme heat, or cools below it, is not known.

In this study, we collected natural *T. oblongifolia* seed from across the species range and combined whole-genome sequencing (WGS), transcriptome analyses, and high-throughput infrared (IR) imaging to resolve a physiological mechanism for extreme heat survival in this species. Seedlings grown at *T*_air_ = 60 °C for 6–8 h daily over at least eight days varied in survival phenotypes. Genome-wide association (GWA) testing identified three “heat tolerance” loci with segregating variation underlying survival differences. IR imaging linked leaf cooling to survival, and RNA-seq implicated pathways for transpirational cooling. Hormone-mediated inhibition of transpirational cooling disrupted survival at 60 °C. Together, we identify genetic variation for survival in extreme heat in *T. oblongifolia* and show that a physiological cooling mechanism enables persistence at the ∼60 °C upper thermal limit of eukaryotic life.

### Natural collections of *T. oblongifolia* seedlings can survive at 60 °C

We first aimed to determine the distribution and natural climates of *T. oblongifolia* to guide the collection of seed for physiology and population genomic investigations of extreme heat tolerance. *T. oblongifolia* clusters in the Mojave Desert and southern Great Basin of the southwestern US, extending into western Arizona and toward the US–Mexico border (*22*, *30*) Rasters of monthly summer maximum temperature (*T*_max_) during 1970-2000 from WorldClim 2 (*31*) overlaid with Global Biodiversity Information Facility (GBIF) occurrences (*30*) and iNaturalist observations (*33*, *34*) revealed that *T. oblongifolia* is restricted to high temperature climates within its species range (Fig. 1A). These data guided our collection of *T. oblongifolia* seed from 5–10 individuals at 40 sites for a total of 223 individuals, categorized into collection regions: “Death Valley” (DV), “Perimeter”, and “Outside”. DV was collected within Death Valley, Perimeter at higher elevations around DV, and Outside at any other site (Fig. 1B; table S1).

**Fig. 1.**
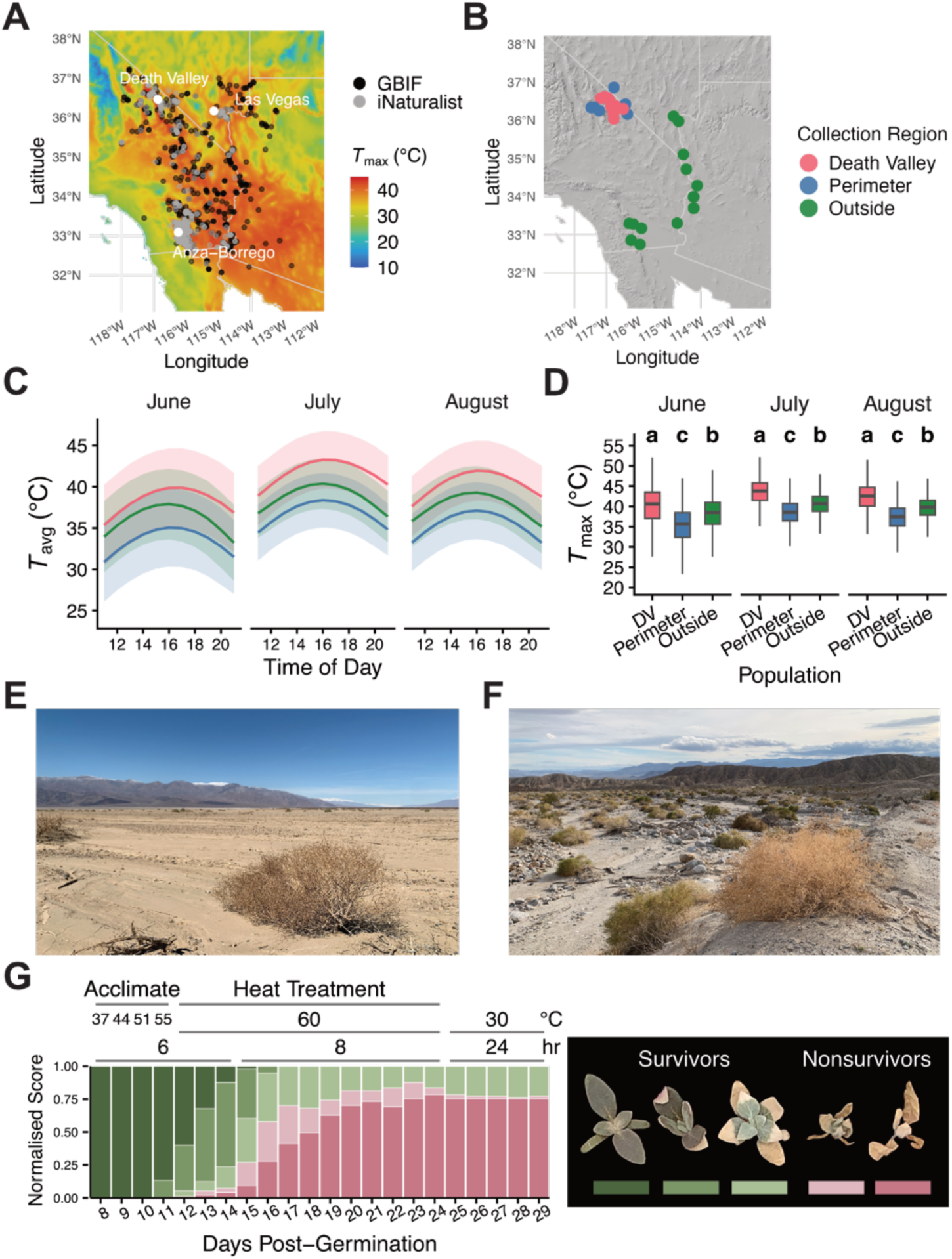
Natural collections of *Tidestromia oblongifolia* seedlings survive at *T*_air_ = 60°C. **(A)** Occurrence records of *T. oblongifolia* from GBIF (*38*, *39*) (black) and iNaturalist observations (*32*) (grey), overlaid on mean summer *T*_max_ (June–August) from WorldClim 2 (*31*). Landmarks Death Valley, Las Vegas, Anza-Borrego in white. **(B)** Seed collection sites, categorized as Death Valley (DV) in pink, Perimeter (higher-elevation sites around DV) in blue, and Outside (all other sites) in green, on a shaded-relief base map. Collection summaries with coordinates in table S1. **(C)** Summer daytime *T*_avg_ (1100–2100) at collection regions by month from ERA5-Land reanalysis (1950–2024; methods) (*33*). Lines are means, shaded bands ± standard deviation (SD). Collection regions differed significantly [linear mixed model, lmer(*T*_avg_ ∼ region × month + hour + (1 | site)); effect of region *F*(_2,37_) = 28.1, *P* = 3.76 × 10^−8^]; pairwise contrasts by estimated marginal means (Tukey-adjusted) in fig. S1. **(D)** Daily *T*_max_ at collection regions by month from the same reanalysis dataset in (C). Letters denote regions that differ significantly [linear mixed model, lmer(*T*_max_ ∼ region × month + (1 | site)); effect of region *F*(_2,37_) = 27.1, *P* = 5.62 × 10^−8^; pairwise contrasts by estimated marginal means, Tukey-adjusted]. DV was hottest, Outside intermediate, and Perimeter coolest across all three months. **(E, F)** Dormant *T. oblongifolia* in the field at Death Valley (E) and Anza-Borrego (F) in February and March 2023, respectively. **(G)** Extreme heat survival assay. Seedlings 8 days post-germination (dpg) were acclimated with stepwise temperature increases (37, 44, 51, 55 °C; 6 h each) followed by heat treatment on 12 dpg at 60 °C (8 h) and recovery at 30 °C continuous. Stacked bars show the normalized distribution of health scores per dpg, corresponding to color below representative survivors (green) and nonsurvivors (pink) shown at right (methods). Growth conditions and daily health status scoring in methods.

Because we grouped collection sites geographically, we next asked whether these regions differed in the temperatures local plants experienced. Hourly summer climate data for 1950–2024 from ERA5-Land Open-Meteo Weather API (*34*) revealed that summer daytime average temperature (*T*_avg_) and *T*_max_ were significantly higher in DV than both Perimeter and Outside, with Outside significantly hotter than Perimeter (*T*_avg_ *P* = 3.76 × 10^−8^; *T*_max_ *P* = 5.62 × 10^−8^) (Fig. 1, C and D, fig. S1A). DV remained consistently hotter across summer months relative to Outside and Perimeter (*P* = 3.8 × 10^−8^) (fig. S1B). Mean summer daytime *T*_avg_ at DV sites reached ∼35–43 °C while Perimeter and Outside reached ∼27–35 °C and ∼30–40 °C, respectively. Absolute summer daytime *T*_max_ at DV sites exceeded ∼50 °C while both Perimeter and Outside had sites with *T*_max_ between ∼45–50 °C.

Our collection included seed from coordinates where the all-time record highest *T*_air_ of 56.7 °C was set in 1913 (*35*) (Fig. 1E). We therefore aimed to determine the upper thermal limit at which *T. oblongifolia* could survive. Seedlings were grown at constant *T*_air_ = 30 °C before sequential daily acclimation to 6 h at 37 °C, 44 °C, 51 °C, and 55.5 °C. The chamber was then set to its maximum *T*_air_ = 60 °C for 6–8 h, repeated for ∼8 days, in an “extreme heat survival assay” (Fig. 1G; methods). Seedlings were scored daily for health status (methods) as “survivors” or “nonsurvivors”, with symptoms of heat-induced stress appearing at *T*_air_ = 55.5 °C and accelerating at *T*_air_ = 60 °C. Overall survival rates varied between experimental rounds (1.96–34.1%). Notably, survivors were still alive and healthy, with newly developed leaf tissue, when treatment at *T*_air_ = 60 °C was extended to 20 days (fig. S2). To disentangle maternal effects from genetic heritability, extreme heat survival was validated in the next generation using seed collected from *T. oblongifolia* plants grown in our greenhouse (fig. S3). These data reveal both a remarkable level of extreme heat survival and phenotypic variation within our natural collections.

### GWA reveals genetic variation underlying survival at 60 °C

As survival phenotypes among *T. oblongifolia* seedlings indicated heritable variation at *T*_air_ = 60 °C, we sought the underlying loci by GWA. Initially, because low-coverage WGS can yield unreliable genotype calls, we generated a species-range reference panel against which genotypes for GWA could be imputed (*36*). One individual per collection region site (fig. S4) was barcoded and sequenced at ∼10× followed by variant calling to identify 3,867,205 single-nucleotide polymorphisms (SNPs) that were phased into reference haplotypes (methods). This panel represented haplotype diversity across our seed collection, providing a reference against which low-coverage genotypes could be resolved.

We next generated samples for association testing, phenotyping our collection for survival at *T*_air_ = 60 °C using our extreme heat survival assay (methods). A total of 1,268 seedlings from 127 individuals (table S2) were assayed in three independent experimental rounds, each with four trays of ∼120 seedlings containing the same genotypes randomized by position. *K*-means clustering of daily health scores resolved survivors from nonsurvivors (methods). The top survivors, the early nonsurvivors, and a random sampling of late nonsurvivors (table S3) were individually barcoded and sequenced at ∼7.5×. Called variants were phased and imputed against the reference panel (methods). Principal component analysis (PCA) showed that survival phenotypes did not separate along the major axes of genome-wide variation (Fig. 2A; methods). These data suggested that SNP associations with survival would not be a consequence of population substructure.

**Fig. 2.**
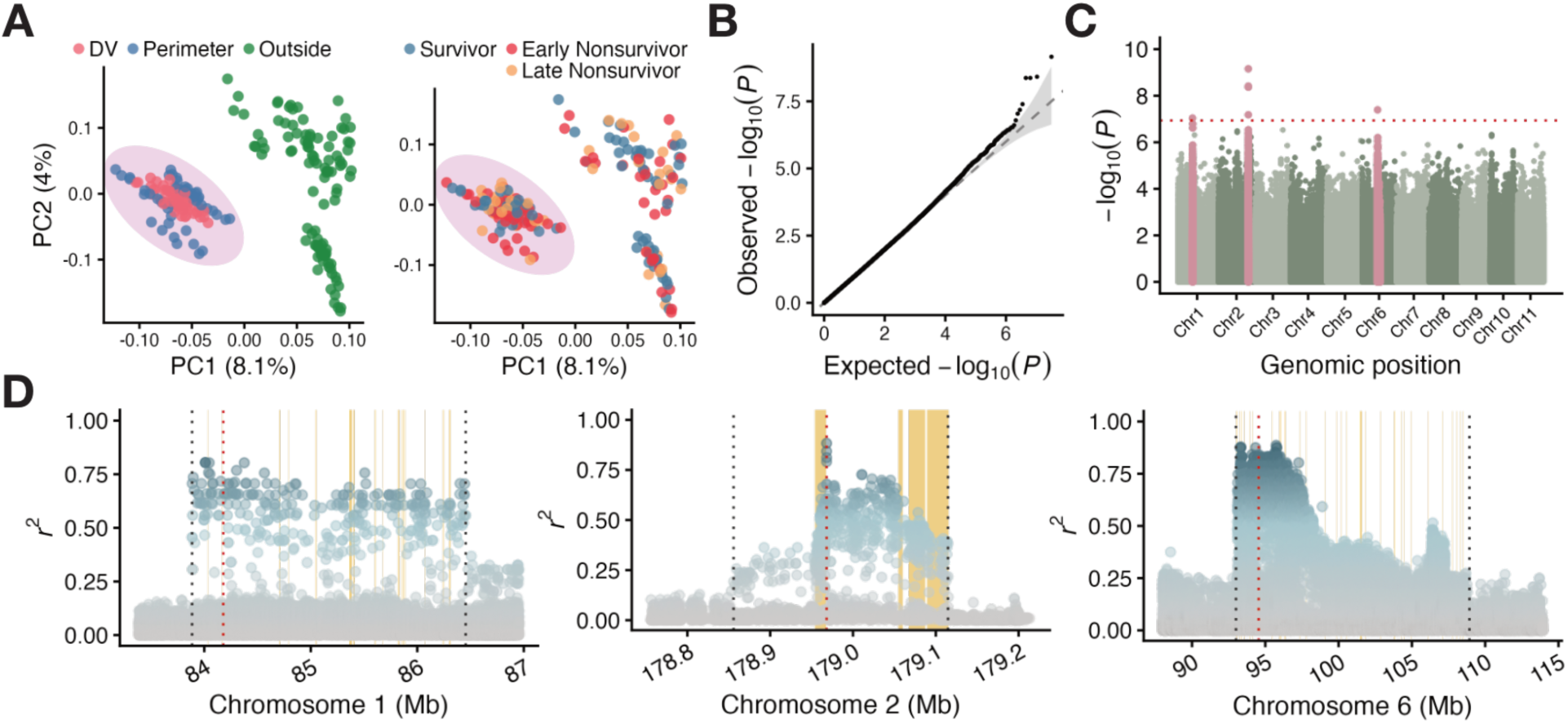
GWA identifies loci associated with extreme heat survival in T. oblongifolia. **(A)** Principal component analysis (PCA) of imputed single-nucleotide polymorphisms (SNPs), linkage-disequilibrium (LD)-pruned in 150 kb windows at squared Pearson correlation coefficient (*r*^2^) < 0.2 (370,948 retained), shown by collection region (left) and survival group (right). Samples separate along principal component (PC) 1 by population structure, with Outside distinct from DV and Perimeter, while survival groups do not separate along the major axes of population structure. Purple ellipse indicates DV and Perimeter samples retained for association mapping (*n* = 107 of *N* = 219 total; table S3). **(B)** Quantile–quantile plot of observed versus expected −log_10_ (*P*); grey band, 95% confidence envelope; diagonal, null expectation. Genomic inflation factor (*λ*_GC_) = 0.983. **(C)** Manhattan plot of association between SNP genotype and relative survival, tested by linear model (methods). Red dotted line, Bonferroni threshold (α = 0.05 over 430,246 LD-pruned variants; −log_10_(*P*) ≈ 6.93). Top SNPs on Chromosome (Chr) 1, Chr 2, and Chr 6 exceed threshold. SNPs within LD boundaries defined in (D) colored in pink. **(D)** LD around the three lead SNPs: Chr 1, 84.18 Mb; Chr 2, 178.97 Mb; Chr 6, 94.55 Mb. Points are SNPs colored by *r*^2^ with the lead variant (red dotted line). Black dotted lines mark LD-block boundaries, set per locus (methods). Vertical yellow lines are candidate gene positions.

To identify loci underlying this variation, we tested each SNP for association with survival. As population structure may generate spurious genotype–phenotype associations, differentiation between regions was first evaluated by PCA and Weir and Cockerham’s estimator of fixation index (*F*_ST_) (*37*, *38*). Outside individuals segregated from clustered DV and Perimeter individuals (Fig. 2A; fig. S5), and *F*_ST_ showed Outside was differentiated from DV and Perimeter regions (table S4). Therefore, association testing was performed with only DV and Perimeter samples (*N* = 107; *n* = 46 early nonsurvivors, 26 late nonsurvivors, and 35 survivors; table S3; methods). From DV and Perimeter only, 15,880,175 SNPs were recovered after imputation. Number of days to death was modeled by Cox proportional hazards regression (*38*, *39*) to derive a continuous “relative survival” phenotype that accommodated individuals that never died. Relative survival phenotypes also did not differ by collection region or site (fig. S6). Each SNP was then tested against relative survival by linear regression (methods). Test statistics showed no evidence of genomic inflation (λGC = 0.983; Fig. 2B), and seven SNPs exceeded a conservative Bonferroni threshold (*P* < 1.162 × 10^−7^; Fig. 2C). Alternate allele dosage for top SNPs is shown against relative survival in fig. S7. After collapsing redundant signal from nearby linked variants, three lead SNPs remained, one each on chromosome (Chr) 1, Chr 2, and Chr 6 (methods). Each lead SNP sat within a region of broadly elevated association signal, bounded by the decay of linkage disequilibrium (LD) with the lead variant (Fig. 2D; methods). We refer to these regions hereafter as “heat tolerance” loci, as these data are consistent with a set of linked variants tagging an underlying causal locus. Together, these results demonstrate segregating genetic variation for survival at *T*_air_ = 60 °C within our natural collection of *T. oblongifolia*.

### Leaf cooling predicts survival at 60 °C

Although *T. oblongifolia* seedlings showed genetic variation for survival at *T*_air_ = 60 °C, it was not clear whether this was through *T*_leaf_ lowering or matching *T*_air_. We developed a high-throughput assay for IR imaging of ∼324 seedlings during extreme heat survival assays (fig. S8; methods). *K*-means clustering of daily health scores grouped individuals by survival phenotypes (fig. S9): cluster 1 = early nonsurvivors, 2 = late nonsurvivors, and 3 = survivors. Kaplan-Meier survival analysis (*25*) confirmed clusters were differentiated for survival probabilities (log-rank *P* < 0.0001; Fig. 3A). On the first day at *T*_air_ = 60 °C, leaf-air temperature offset (Δ*T* = *T*_leaf_ − *T*_air_) differed significantly among clusters, with survivors recording the most negative Δ*T* over time. Δ*T* scaled with days survived: the late nonsurvivors recorded a more negative Δ*T* than the early nonsurvivors (*P*(cluster) = 2.66 × 10^−5^; Fig. 3, B and C). A gradual change to a less negative Δ*T* was observed over time—likely the result of insufficient chamber dehumidification (fig. S10). A Bayesian generalized linear mixed model indicated that the only credible predictor of mortality was higher *T*_leaf_ (pMCMC < 0.001; Fig. 3D). Although tray position weakly predicted *T*_leaf_, neither leaf morphology nor positional effects predicted mortality when *T*_leaf_ was excluded (fig. S11, A and B), and energy balance modeling suggested leaf area and shape were too uniform to impact Δ*T* (fig. S11C). These data reveal that *T. oblongifolia* does lower *T*_leaf_ below *T*_air_ to survive at 60 °C, and that magnitude of Δ*T* can predict survival in extreme heat.

**Fig. 3.**
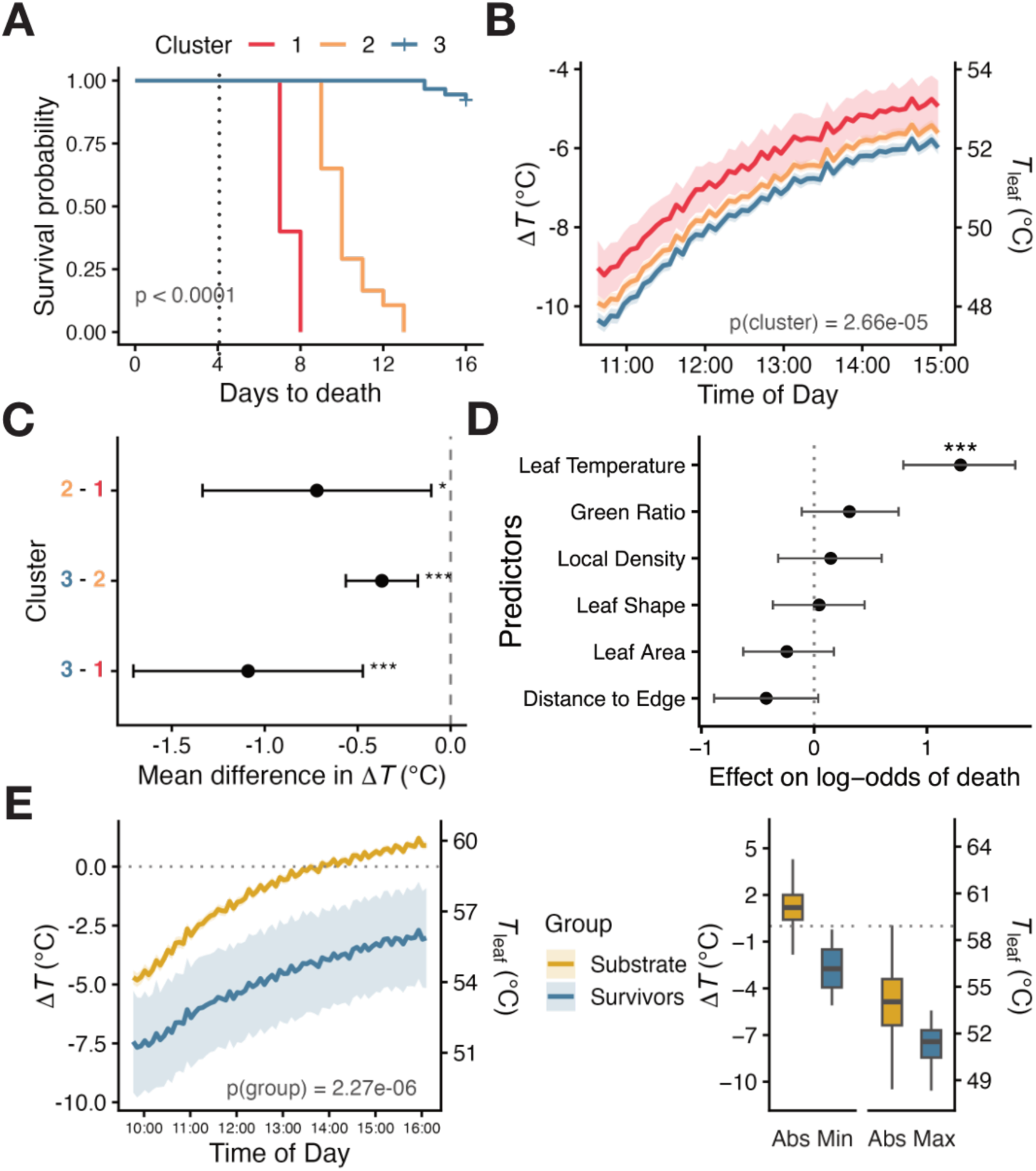
Leaf cooling predicts *T. oblongifolia* survival at *T*_air_ = 60 °C. **(A)** Kaplan–Meier survival curves (*38*, *39*) for plants partitioned into three clusters by *k*-means clustering (k = 3) of normalized daily health-score trajectories (methods). Clusters differed in survival (log-rank *P* < 0.0001). Dotted line indicates start of *T*_air_ = 60 °C. *n* = 5, 103, 90 for clusters 1, 2, 3, repsectively. **(B)** Leaf-air temperature offset (Δ*T* = *T*_leaf_ − *T*_air_; right axis, *T*_leaf_) through *T*_air_ = 60 °C by cluster. *n* as in (A). Lines are means, shaded bands 95% confidence interval (CI). Clusters differed significantly [linear mixed model, lmer(Δ*T* ∼ cluster × time + (1 | cell)); *P*(cluster) = 2.66 × 10^−5^]. **(C)** Pairwise differences in mean Δ*T* between clusters, estimated marginal means from (B), unadjusted; points, mean difference (°C); bars, 95% CI. \**P* < 0.05, \*\**P* < 0.01, \*\*\**P* < 0.001. **(D)** Effect of standardized leaf traits on log-odds of death (Bayesian generalized linear mixed model, MCMCglmm, categorical; tray and individual as random effects; *n* = 197). Estimates per 1 SD; points, posterior means; bars, 95% credible intervals. *T*_leaf_ was the only credible predictor (***pMCMC < 0.001). **(E)** Left: Day 8 of *T*_air_ = 60 °C. Δ*T* for survivors versus substrate; plotted as in (B) [lmer; *P*(group) = 2.27 × 10^−6^]. Right: Absolute minimum and maximum Δ*T* per plant. Boxes, median and interquartile (IQR) range; whiskers, 1.5 × IQR. Groups differ significantly (one-way analysis of variance (ANOVA); minimum, *P* = 0.012; maximum, *P* = 6.61 × 10^−9^). *n* = 4 survivors, *n* = 229 substrate cells.

We next investigated whether *T. oblongifolia* survivors maintained a strong negative Δ*T* after extended days at *T*_air_ = 60 °C. Indeed, on the eighth day of heat treatment, the absolute maximum negative Δ*T* of survivors was −10.57 °C (*P* = 6.61 × 10^−9^; Fig. 3E), comparable to Δ*T* observed on the first day at *T*_air_ = 60 °C (Fig. 3B). Astonishingly, we also observed that *T*_leaf_ of survivors ranged from 53.54 °C to 58.76 °C in the last 30 min of heat treatment. The eukaryotic upper thermal limit is known to be ∼60 °C (*4–7*). Taken together, these data indicate that *T. oblongifolia* seedlings maintain a lower *T*_leaf_ than *T*_air_ over extended days at 60 °C, but are also adapted to tolerate some exposure to *T*_leaf_ matching *T*_air_ at the upper thermal limit of eukaryotic life.

### Transcriptional variation underlies leaf cooling phenotypes

IR imaging showed that survivors cooled more than nonsurvivors (Fig. 3, B and C), but not whether this difference had a molecular basis. We therefore asked whether leaves differing in Δ*T* also differed transcriptionally. RNA-seq reads were aligned to the *T. oblongifolia* AmA10 reference assembly (*41*) and differential expression (DE) was tested with DESeq2 (*42*, *43*) (methods). Of 30,294 annotated genes, 20,967 passed filtering, of which ∼50% (9209) were differentially expressed genes (DEGs). We analyzed these data in two ways: the overall transcriptional response across heat acclimation, and the direct contrast between differently cooling leaves.

To place the cooling-associated signal in the context of the overall heat response, we first evaluated transcriptome changes across heat acclimation. Leaves were sampled during timepoints at *T*_air_ = 30 °C, 37 °C, 44 °C, 51 °C, and from the lowest (55_cold) and highest (55_hot) *T*_leaf_ individuals at *T*_air_ = 55.5 °C. Developmental controls at *T*_air_ = 30 °C were sampled in parallel each day (d9–d13; fig. S12A). Samples were not taken at *T*_air_ = 60 °C to avoid confounding cell-death signatures with the heat response. Relative transcriptional changes to heat were evaluated by comparing each timepoint back to the baseline (*T*_air_ = 30 °C), after controlling for development. PCA on variance-stabilized counts separated 51 °C and 55.5 °C samples from all others along PC1, accounting for 59.6% of total variation. 55_cold and 55_hot samples segregated along PC2, indicating unique responses in differently cooling leaves (Fig. 4A). Most DEGs were shared between all samples except for 55_hot with ∼55% of its 5158 DEGs unique from any other sample (fig. S12, B and C). These data suggest a strong global transcriptional response to extreme heat in *T. oblongifolia* that results in variation in leaf cooling phenotypes.

**Fig. 4.**
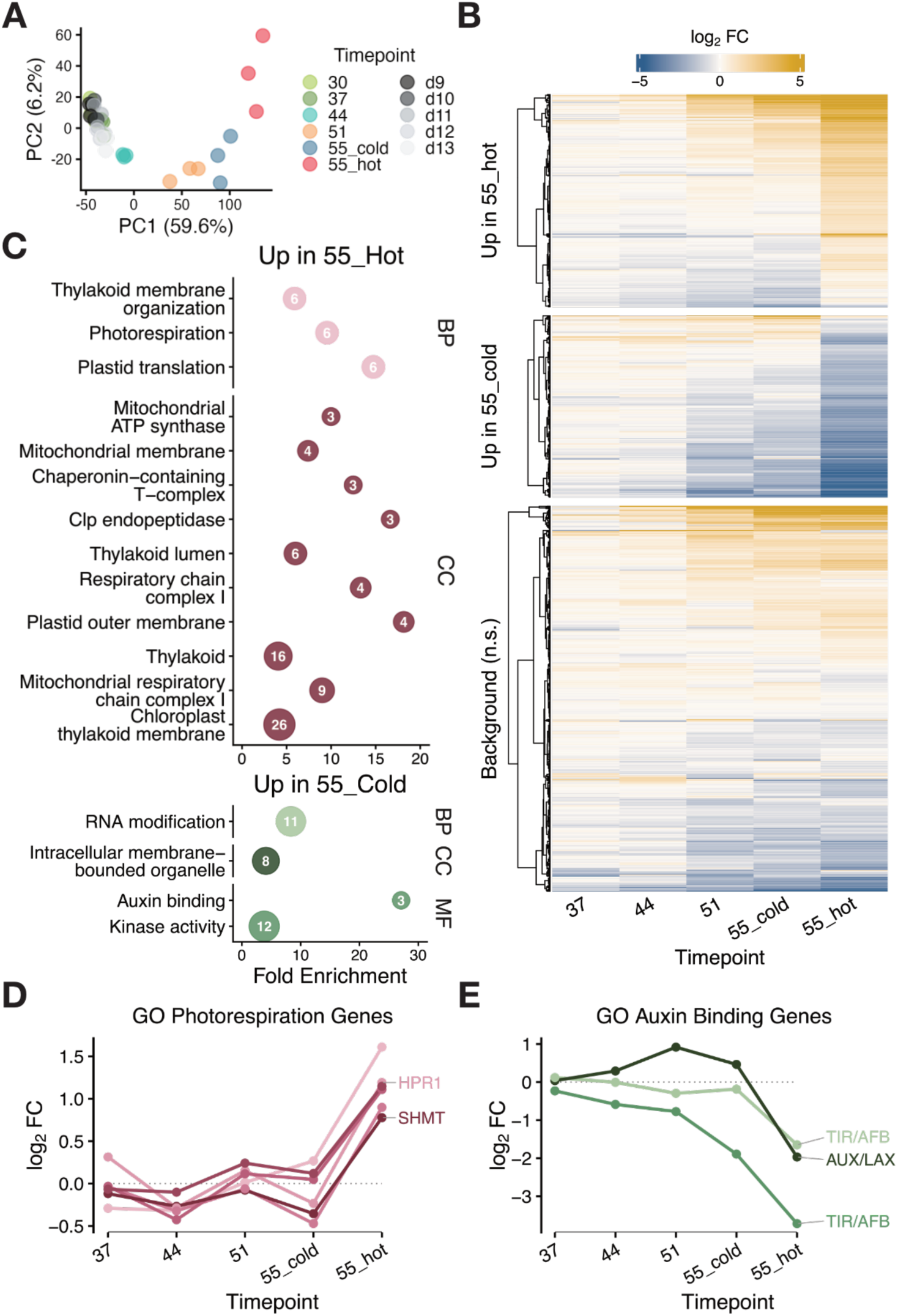
RNA-seq reveals molecular signatures of transpirational cooling in T. oblongifolia. **(A)** PCA of variance-stabilized counts across heat-treatment timepoints (30, 37, 44, 51 °C; 55_cold = coolest 55 °C leaves, 55_hot = warmest 55 °C leaves) and day-matched controls sampled at 9–13 dpg (d9–d13, matched to 30, 37, 44, 51, and both 55 °C groups respectively; *n* = 3 per timepoint) (fig. S12A). PC1 and PC2 explain 59.6% and 6.2% of variance. **(B)** Differentially expressed genes (DEGs) between 55_cold and 55_hot (apeglm-shrunk; s-value < 0.005, |log_2_ fold change| > log_2_ 1.5), split into “up in 55_hot”, “up in 55_cold”, and random set of 600 non-significant background genes. Values are log_2_ fold change (L2FC) of each timepoint against its day-matched control, minus the same contrast at baseline (30 °C). Rows clustered by Ward’s D2; columns ordered by timepoint. **(C)** GO term enrichment among genes up in 55_hot (top) and up in 55_cold (bottom) in (B), by biological process (BP), cellular component (CC), and molecular function (MF). Point position, fold enrichment; number in point, genes in term. Terms shown at q < 0.05 with ≥ 3 genes (hypergeometric test, Benjamini–Hochberg corrected). **(D)** L2FC as in (B) across developmentally controlled timepoints, relative to developmentally controlled baseline, for genes in the photorespiration GO term (enriched among genes up in 55_hot) in (C). Canonical photorespiration homologs HPR1 and SHMT are labelled. **(E)** L2FC as in (B) in the auxin binding GO term (enriched among genes up in 55_cold) in (C). Labelled by Arabidopsis ortholog family (TIR/AFB, AUX/LAX).

As *T*_leaf_ was predictive of survival (Fig. 3D), transcriptomics signatures of 55_cold and 55_hot samples provided a particularly useful contrast to identify molecular mechanisms for survival in extreme heat. Direct comparison of 55_cold and 55_hot samples revealed 614 DEGs (Fig. 4B), ∼30% of which were unique from any other sample’s DEGs (fig. S12, B and C). Gene Ontology (GO) enrichment (*44*) revealed unique signatures in 55_cold and 55_hot transcriptomes (Fig. 4C). Notably, photorespiration was an enriched GO term in the 55_hot samples (Fig. 4C, above), and is a signature of stomatal closure and therefore reduced transpirational cooling (*44*, *45*). Expression patterns of GO enriched photorespiration genes indicated that all six were upregulated in 55_hot samples versus 55_cold (Fig. 4D, left), with two *T. oblongifolia* genes homologous to core photorespiration genes. Independent evaluation of other core homologous photorespiration genes also showed upregulation in 55_hot samples compared to cold (fig. S13A). In contrast, auxin binding was a significantly enriched GO term in 55_cold samples (Fig. 4C, below), with core auxin binding genes downregulated in 55_hot samples compared to 55_cold (Fig. 4D, right; fig. S13B). Auxin binding is known to directly induce stomatal opening, increasing transpirational cooling rates (*45*, *46*). These data show a clear transcriptome difference in the cellular states of hot and cold leaves at 55 °C, suggesting molecular mechanisms for physiological cooling differentiate *T. oblongifolia* survivors from nonsurvivors in extreme heat.

### Leaf cooling enables survival at 60 °C

As transcriptomics suggested molecular evidence for extreme heat survival through transpirational cooling, we investigated if cooling was adaptive and tested if inhibition of cooling would disrupt survival. Δ*T* was measured over changing *T*_air_ = 51 °C, 55 °C, and 60 °C (Fig. 5A). Absolute minimum and maximum Δ*T* of survivor cluster 3 were significantly lower than nonsurvivor clusters 1 and 2 at *T*_air_ = 60 °C and 55.5 °C (*P* < 0.005), but not 51 °C (*P* > 0.2) (Fig. 5A, left). The slope of the relationship between Δ*T* and *T*_air_ (β) was calculated across 51 °C, 55.5 °C, and 60 °C (*46*). β was negative for all clusters in both absolute minimum and maximum slopes, with survivor cluster 3 significantly more negative than late nonsurvivor cluster 2 (*P* = 0.0083; *P* = 0.017) (Fig. 5A, right). Energy budget modeling was used to determine whether observed Δ*T* was theoretically plausible. Modeling chamber parameters with varying stomatal conductance (*g_sw_*) showed that a Δ*T* between −10 °C to −13 °C when *T*_air_ = 60 °C could be expected (Fig. 5B). Indeed, an absolute maximum negative Δ*T* of −13.03 °C was observed (Fig 5A), indicating that *T. oblongifolia* cools to the extreme of its physiological limit at *T*_air_ = 60 °C. These data indicate *T. oblongifolia* survivors have an increased capacity for adaptive leaf cooling over nonsurvivors.

**Fig. 5.**
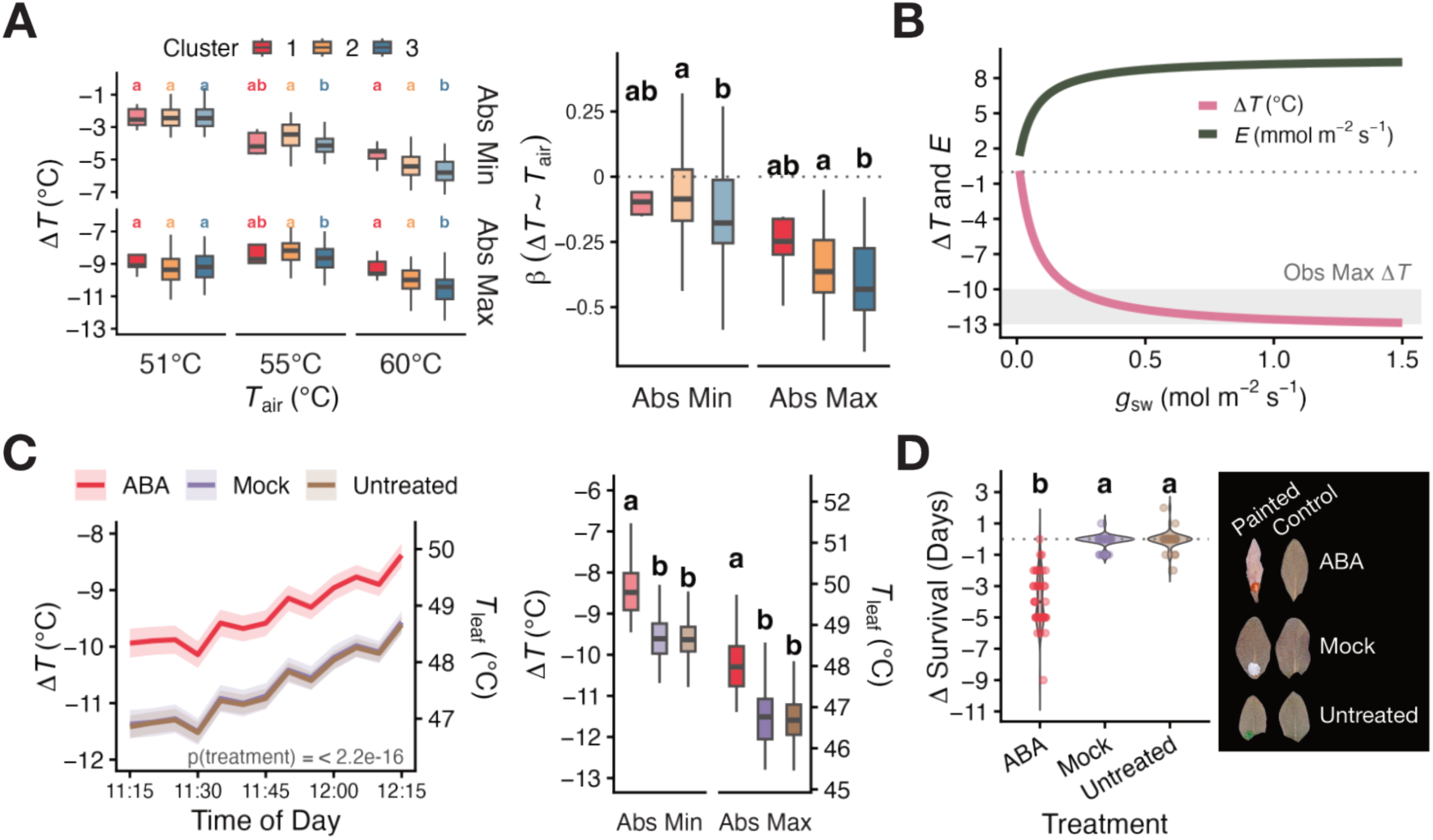
ABA-mediated inhibition of cooling accelerates time to death at *T*_air_ = 60 °C. **(A)** Left: absolute minimum and maximum Δ*T* per plant by cluster at *T*_air_ = 51, 55, and 60 °C, from the same imaging data and cluster assignments as Fig. 3A–D. Boxes, median and IQR; whiskers, 1.5 × IQR; letters denote clusters differing within each panel (ANOVA with Tukey honestly significant difference (HSD); clusters differed at 55 °C and 60 °C, *P* < 0.005, but not at 51 °C, *P* > 0.2). Right: per-plant slope of Δ*T* on *T*_air_ (β), by cluster (ANOVA with Tukey HSD; absolute minimum, *P* = 0.0083; absolute maximum, *P* = 0.017). **(B)** Modeled Δ*T* and transpiration rate (*E*) against varying stomatal conductance (*g_sw_*), from steady-state leaf energy balance (methods). Grey band represents observed maximum Δ*T* range. Dotted line, Δ*T* = 0. **(C)** Left: Δ*T* (right axis, *T*_leaf_) through *T*_air_ = 60 °C. Lines are means, bands 95% CI. Treatments differed significantly [lmer(Δ*T* ∼ treatment × time + (1 | cell)); *P*(treatment) < 2.2 × 10^−16^]. Right: absolute minimum and maximum Δ*T* per leaf; letters denote treatments differing (ANOVA with Tukey HSD; minimum and maximum *P* < 2 × 10^−16^). *n* = 41 ABA, *n* = 40 mock, *n* = 40 untreated. Mock and Untreated datapoints overlap. **(D)** ΔSurvival (days to death of painted leaf − internal control leaf). Violins, distribution; boxes, median and IQR; points, individual plants. Letters denote treatments differing (ANOVA with Tukey HSD; *P* < 2 × 10^−16^). *n* = as in (C). Negative values indicate the painted leaf died first. Right, representative leaves.

To test whether leaf cooling actually impacts survival in extreme heat, we used abscisic acid (ABA) to induce stomatal closure to inhibit transpirational cooling in *T*_air_ = 60 °C (*47*). Leaves were painted with 50 µM ABA, mock (DMSO 0.05%), or left untreated ∼20 min before heat treatment at 60 °C. A significantly less negative Δ*T* was immediately recorded in the ABA-treated leaves compared to mock or untreated (*P*(treatment) < 2.2 × 10^−16^) (Fig. 5C). ABA, mock, or untreated leaves were scored daily for survival compared to an internal control leaf of the same developmental stage on the same plant. ABA-treated leaves showed an accelerated time to death compared to the internal control leaf, dying on average four days sooner, while mock-treated and untreated leaves were not different from their controls (*P* < 2 × 10^−16^) (Fig. 5D). These data show that transpirational cooling is a key mechanism for *T. oblongifolia* survival at the ∼60 °C upper thermal limit of eukaryotic life.

## Discussion

*T. oblongifolia* is a unique organism that has adapted to an extreme environment nearly incompatible with eukaryotic life. The molecular mechanisms of less complex thermophilic eukaryotes, such as fungi and unicellular algae, are well documented (*7*, *8*, *47–51*). By contrast, we find that this complex, multicellular plant survives at the ∼60 °C upper thermal limit of eukaryotic life through a physiological cooling mechanism.

Using seedlings from our collection of *T. oblongifolia* seeds across the species range (Fig. 1B), IR imaging revealed leaf cooling correlated with survival during maximum thermal limit testing at *T*_air_ = 60 °C (Fig. 3). Comparing transcriptomic signatures of differently cooling leaves revealed signals on genes and pathways that indirectly regulate transpiration (Fig. 4). We tested the role of cooling in survival through ABA-mediated inhibition of stomatal transpiration at 60 °C, which resulted in a less negative Δ*T* by about 1–1.5 °C (Fig. 5C) and an accelerated time to death (Fig. 5D). Together, these data suggest that transpirational cooling is a key component of extreme heat survival in *T. oblongifolia*.

GWA testing supported a genetic basis for variation in extreme heat survival and identified three heat tolerance loci. However, resolving causal variants and genes at these loci will require follow-up fine mapping and functional validation. Our transcriptomic data offer an independent route to identifying gene candidates, such as the 20 genes at heat tolerance loci that were differentially expressed in our heat response assays (fig. S12, D and E). This is a promising pool of priority candidates for follow-up investigations. Yet, this positional overlap does not capture causal variants acting in *trans*, nor those altering protein function rather than expression, and further refinement of signal at heat tolerance loci would provide a starting point for tracing such effects. Our data nonetheless reveal high-priority gene candidates that can immediately be assayed for function in extreme heat survival.

Prior to this study, we did not know if *T. oblongifolia* survived high-temperature environments through leaf cooling or through an adaptation to *T*_leaf_ matching *T*_air_. Our data show leaf cooling as a critical component enabling survival; however, we did also observe *T*_leaf_ of survivors at ∼54– 59 °C after at least eight days (Fig. 3E), and as long as 20 days (fig. S2), at *T*_air_ = 60 °C. Both physiological and molecular adaptations therefore appear to enable survival at extreme high temperatures: conceivably, thermostability conferred through amino acid substitutions in proteins and enzymes, and changes in membrane or cell wall composition. Additionally, over those 8–20 days at *T*_air_ = 60 °C, seedlings grew new, healthy leaf tissue (fig. S2). Although *T. oblongifolia* growth is optimal at 44 °C (*22*), this suggests that developmental pathways are thermostable in survivors. The molecular properties of extreme heat adaptation in *T. oblongifolia*, beyond the extreme leaf cooling determined here, remain an open and enticing questions.

Many plants use cooling mechanisms to tolerate increasing temperatures (*52*), some even transpiring to their own detriment (*19*, *53*), but none are known to cool to Δ*T* = −13 °C at *T*_air_ = 60 °C (Fig. 3 & 5). How *T. oblongifolia* does so without catastrophic water loss is unclear, but part of the answer may lie in water supply and transport: Björkman and colleagues recorded low stomatal resistance *in situ* in Death Valley (*50*, *51*), and low whole-plant hydraulic resistance in controlled conditions (*52*). They concluded that *T. oblongifolia* must have access to water where it occupies Death Valley (*54*). However, water supply and transport alone cannot explain survival at 60 °C as unrestricted water would not confer comparable heat tolerance on most species. Furthermore, *T. oblongifolia* nonsurvivors did cool, just not as deeply and with a weaker response than survivors as *T*_air_ rose (Fig. 3, B and C; Fig. 5A). This variation in the magnitude of cooling could reflect a cost that survivors are better able to bear, with season-to-season variability in upper thermal limits maintaining that variation in the population. Understanding not only how *T. oblongifolia* cools better than other species, but also how survivors cool better than nonsurvivors, presents a promising opportunity to fully understand a physiological trait shared across plants.

Whether discovery of heat tolerance strategies in wild plants will provide real solutions for crop improvement is still undetermined (*55*, *56*). Global warming patterns are increasingly evident and threaten crop systems and agricultural practices (*57*). In this study, we have identified two potential routes to uncovering novel strategies for crop improvement or engineering. If we can understand the strategies *T. oblongifolia* has adapted for physiological cooling, a shared trait among plants, it may be possible to increase upper thermal limits in some crops. Likewise, if we can uncover the molecular mechanisms of adaptation to the upper thermal limit of eukaryotic life, we may be able to generate enhanced molecular thermostability in crops. An increase of upper thermal limits of only a few degrees in crops could substantially buffer future climate-driven yield losses to increasing temperatures (*31*).

## Acknowledgements

We thank Dr Marco D’Ario for early input and assistance with field collections; Drs Laura Leventhal, Flavia Bossi, Evan Saldivar, Karine Prado, Colette Berg, and Xing Wang for input and feedback; Professors Dmitri Petrov and Moi Exposito-Alonso for early feedback on project design; Anna Somerville for horticulture support; Drs Matt Stata, Shifeng Cheng, and Hongbing Liu for *Tidestromia oblongifolia* genomics resources; Carnegie Science Department of Plant Biology facilities team: Ismael Villa, Johnny Materassi-Shultz, Angelica Vazquez, and Theo Van De Sande for maintaining equipment and materials. Claude Sonnet 4.6, Sonnet 5, and Opus 5 (Anthropic) were used to assist with drafting and editing portions of the manuscript text, and all content was reviewed and approved by the authors, who take full responsibility for it. This work was supported in part through computational resources and services provided by the Institute for Cyber-Enabled Research at Michigan State University. Lastly, we are grateful to Carnegie Science for having supported desert plant research, which remains a deeply underexplored field, since 1903.

## Funding

Carnegie Canada Grant (JMF).

Plant Resilience Institute Postdoctoral Fellowship (JMF).

Carnegie Venture Program (SYR).

The U.S. National Science Foundation grant DBI-2419923 (SYR).

The U.S. National Science Foundation grant IOS-2312181 (SYR).

The U.S. National Science Foundation grant IOS-2406533 (SYR).

The U.S. National Science Foundation grant OISE-2434687) (SYR).

U.S. Department of Energy, Office of Science, Office of Biological and Environmental Research, Genomic Science Program grant DE-SC0021286 (SYR).

U.S. Department of Energy, Office of Science, Office of Biological and Environmental Research, Genomic Science Program grant DE-SC0021286 DE-SC0023160) (SYR).

## Permits

Death Valley National Park (DEVA-2022-SCI-0020).

Anza-Borrego Desert State Park (CDD-2023-008-ABDSP).

## Author Contributions

Conceptualization: JMF, MCB, SYR.

Methodology: JMF, MCB, TDS, DBL, SYR.

Investigation: JMF.

Formal Analysis: JMF.

Visualization: JMF.

Funding Acquisition: JMF, SYR.

Writing (original draft): JMF.

Writing (review and editing): JMF, MCB, SYR, DBL, TDS.

## Data, code, and materials availability

## Supplementary Materials

Materials and Methods.

Figs. S1 to S13.

Tables S1 to S4.

References (1–62).

Data S1.

