## Supplementary material for "Extreme cooling enables survival in extreme heat": materials and methods

Materials & Methods

**Plant material and growth**

***Extreme heat "Death Valley" cabinet.*** Extreme heat experiments were carried out using a customized Bronson Climate Incrementum 3000 plant growth cabinet, built for our extreme heat assays to be conducted at 60 °C, 30% relative humidity (RH), and ~1200 Photosynthetic Photon Flux Density (PPFD).

***Substrate for Tidestromia oblongifolia seedlings.*** Substrate composed of 1:2 ratio of PRO-MIX® HP Mycorrhizae to Turface Athletics MVP clay with ½ scoop Osmocote 14-14-14.

***Germination of Tidestromia oblongifolia seed.*** Seed was germinated in 90 mm petri dish with filter paper and ~5 mL distilled Milli-Q® water for 24 h at 14 h photoperiod, ~100–150 PPFD, 30 °C constant, and 50% RH in Death Valley cabinet (*1*).

***Threshing Tidestromia oblongifolia seed.*** Wild seed was threshed through a metal colander with micro-perforated holes (2–3 mm), winnowed with a fan on low speed, and stored in glassine bags on silica beads within a lock-lid container at room temperature.

***Standard growth conditions of Tidestromia oblongifolia seedlings.*** Seedlings were transplanted from germination plates (described above) to Proptek 231 deep cell seed starting tray filled with substrate described above. Seedlings were grown at 14 h photoperiod, 30 °C constant, ~100–150 PPFD, and 50% RH for 48 h in Death Valley cabinet. Cabinet settings were then adjusted to ~1200 PPFD and 30% RH continuously onwards.

***Greenhouse propagation of collected Tidestromia oblongifolia seed.*** Seed was germinated as described above and transplanted to 3" × 3" nursery pots containing substrate described above for ~3 weeks at 14 h photoperiod, ~100–150 PPFD, 30 °C constant, and ambient RH. Seedlings were then transplanted to 8" × 3.5" × 3" containing substrate described above for ~12 months in a greenhouse set to 14 h photoperiod, 30 °C, ~100–150 PPFD, and ambient RH. Seed was collected from branches by suction with shop-vac and threshed as described above.

***Seedling genotypes.*** Seedlings from across the species range were used for genome-wide association (GWA) testing (Fig. 2; table S1 and S2) and for infrared (IR) analysis of differential leaf cooling (Fig. 3). To limit genetic variation among individuals for transcriptomics (Fig. 4) and abscisic acid (ABA) experiments (Fig. 5), and as no site yielded sufficient seed alone, seed for these experiments came from four Perimeter sites that clustered together in principal components analysis (PCA) (sites 1, 13, 14, 15; fig. S4B; fig. S5C).

**Species distribution and site climate**

***T. oblongifolia occurrence data and climate extraction.*** *Tidestromia oblongifolia* occurrence records were obtained from the Global Biodiversity Information Facility (GBIF) (*2*) via the rgbif package (*3*, *4*) and from iNaturalist research-grade observations via rinat v0.1.10 (*5*), retaining only georeferenced records (Fig. 1A). Spatial outliers were removed from each source separately by Mahalanobis distance, excluding records beyond the $\chi^2$ 0.95 (GBIF) and 0.999 (iNaturalist) quantiles (df = 2). Maximum temperature ($T_{max}$) layers were obtained from WorldClim 2 (*6*) at 2.5-arc-minute resolution using geodata v0.6.9 (*7*), and mean summer $T_{max}$ was calculated across June, July, and August. GBIF and iNaturalist Occurrence records were plotted over this layer. Maps were produced in R v4.5.2 using sf v1.1.0 and terra v1.9.11 (*8*, *9*).

***Collection region site climate data and analysis.*** Hourly climate data for June, July, and August during 1950–2024 at each regional collection site were obtained from the ERA5-Land reanalysis (*10*) via the Open-Meteo API (*11*), with daytime taken as hours 1100–2100 (Fig. 1, C and D). Differences in summer daytime average temperature ($T_{avg}$) and $T_{max}$ among site groups (Death Valley, DV; Perimeter; Outside) were tested by linear mixed model with category, month, and hour as fixed effects and site as a random effect (lme4 v2.0.1) (*12*), (lmerTest v3.2.1) (*13*), with pairwise contrasts by estimated marginal means (emmeans v2.0.2) (*14*) (fig. S1). Diel temperature exposure was summarized per site as the daytime area under the temperature curve by trapezoidal integration (pracma v2.4.6) (*15*) and compared among categories by one-way analysis of variance (ANOVA) with Tukey Honestly Significant Difference (HSD).

***Field collections of Tidestromia oblongifolia.*** Field collections at 40 sites across the species range were carried out late February in the DV region (28 sites) and Early March in southern California (12 sites) in 2023 (Fig. 1, B, E and F). Approximately 50–100 seed were collected from 5–10 individuals (223 total) per site from distal branches. Refer to table S1 for exact counts, identifiers, and GPS coordinates.

**Extreme heat survival assay and phenotyping**

***Extreme heat assay of Tidestromia oblongifolia seedlings.*** Seedlings for extreme heat survival assays were germinated and sown in standard conditions as described above. Additionally, trays were taped along cell boundaries with industrial grade ½ inch duct tape (GGR Supplies; CDT-36) and edges were saran-wrapped with bottom-tray to reduce evaporation. Initial transplanting was performed 24 h after germination between 0900 and 1300, with supplementary transplanting 48–72 h after germination. Each experimental round comprised four trays containing the same genotypes, randomized by position within each tray. Of 162 seedlings transplanted per tray, ~25% failed to establish and died before leaf-tissue harvesting at 9 days post-germination (dpg), leaving ~120 seedlings per tray. Each round aimed for approximately $N$ = 8 technical replicates of one genotype (sibling seed from the same parent plant), with approximately $n$ = 2 per tray. Approximately 30 min before heat treatment each day, trays were bottom-watered until the substrate was in contact with water and rotated through chamber positions on a fixed schedule. Plants were not water-limited to replicate *in situ* environments (*16*, *17*). From 12 dpg, seedlings were heat acclimated under baseline growth conditions (~1200 PPFD, ~30% RH, 14 h photoperiod, 30 °C constant), with a 6 h heat treatment programmed mid-photoperiod and increasing daily from 37 to 44, 51, and 55 °C. Thereafter the chamber maximum of 60 °C was applied for the remainder of the experiment (Fig. 1G), initially for 6 h, extended to 7 h if no change in health score was observed the following day, and to 8 h if no change was observed at 7 h. Once the number of survivors and nonsurvivors did not change for three days, the experiment was terminated. If experiment was run without tissue sampling at 9dpg (fig. S2), heat acclimation was initiated on 8 dpg (Fig. 1G).

***Chamber environmental monitoring.*** Air temperature ($T_{air}$) and RH were logged inside the Death Valley cabinet at 5 min intervals with an Onset® HOBO® U12-012 data logger, centered to the left or right of trays at leaf-surface level. Measured $T_{air}$ was adjusted for excess radiation heat absorbed by device.

***Health status scoring.*** White-light images were taken ~30 min before heat acclimation or heat treatment (fig. S8B). Seedlings in images were assigned one of five categories of visible heat damage: 0 (dark green) = no visible damage, 1 (green) = minor leaf damage, 2 (light green) = moderate leaf damage, 3 (light pink) = severe leaf damage, 4 (dark pink) = dead. Scores of 1–2 reflected increasing leaf damage in otherwise healthy plants; scores of 3–4 reflect damage from which plants did not recover. All images from one experiment were scored in a single session.

Seedlings were scored from the day before heat acclimation until the number of survivors and nonsurvivors did not change for three consecutive days. A score of four was recorded as death and used as the survival endpoint. Scores were normalized per tray.

***Clustering of health-score trajectories.*** Seedlings were grouped by daily health status scores described above. Seedlings that died before heat acclimation were excluded from downstream analysis. A terminal-day score of 4 was appended so that survivors had a defined time-to-death event. Scores of 1–3 were re-recoded to 0, giving a binary alive/dead series per seedling. The series were centered and scaled per day, and seedlings clustered by *k*-means (nstart = 10, seed = 42), with the number of clusters chosen from the total within-cluster sum of squares across increasing *k*. Analyses were performed in R.

***Kaplan–Meier survival analysis.*** Survival curves were estimated by the Kaplan–Meier method (survival v3.8.6) (*18*, *19*) and compared between clusters by log-rank test, then plotted with survminer v0.5.2 (*20*). Time to death was the number of days until the first score of 4, with surviving seedlings censored at the end of the experiment. Analyses were performed in R.

**Whole-genome sequencing: design and sampling**

***Study design.*** Two independent genome-wide sequencing datasets were generated, 1) a reference panel from representatives across collection regions for downstream imputation (*N* = 38; fig. S4), and 2) a survival panel for GWA testing with relative survival phenotypes (*N* = 225) (Fig. 2A; fig. S5A).

***Genomic DNA extractions from whole leaves.*** One leaf was harvested 9 dpg, flash frozen in liquid nitrogen, and stored at −80 °C. Tissue was homogenized in 2 mL Eppendorf® tube with one 3 mm Tungsten Carbide ball in Geno/Grinder® for two rounds of 1 min at 25 strokes per second, re-frozen in between. Approximately 10 mg of homogenized tissue was lysed in 500 µL 2× cetyltrimethylammonium bromide (CTAB) buffer (1.4 M NaCl, 100 mM Tris-HCl pH 8.0, 20 mM EDTA pH 8.0, 2% w/v CTAB, 10% w/v PVP-10, with 5% v/v 2-mercaptoethanol added immediately before use) preheated to 65 °C. Lysates were incubated at 65 °C for 50 min with periodic inversion, cooled at room temperature for 10 min, and extracted with an equal volume of chloroform:isoamyl alcohol (24:1). Following centrifugation at 20,000 rcf for 20 min at 20 °C, the aqueous phase was precipitated with 0.75 volumes of room-temperature isopropanol and pelleted at 20,000 rcf for 15 min at 20 °C. Pellets were washed twice in 70% ethanol, air-dried, and resuspended in 50 µL water + 20 µg/mL RNase A. Genomic DNA was evaluated for purity by Nanodrop (A260/A280 and A260/A230 ratios), quantified by Qubit, and stored at −80 °C.

**Whole-genome sequencing: reference panel**

***Reference panel: whole-genome sequencing.*** Libraries were prepared by the Texas A&M AgriLife Genomics and Bioinformatics Service with the PerkinElmer NEXTFLEX Rapid XP DNA-Seq Kit HT and sequenced by Novogene on an Illumina NovaSeq X Plus (PE150) to a target depth of ~10× ($N$ = 38). Realized coverage after duplicate marking averaged 8.1× (median 6.9×, range 3.6–18.4×).

***Reference panel: read processing and alignment.*** Read integrity was verified by MD5 checksum, read quality assessed with FastQC v0.12.1 (*21*) and aggregated with MultiQC v1.14 (*22*) before and after trimming. Reads were trimmed with fastp v0.24.0 (*23*) with paired-end adapter auto-detection and poly-G and poly-X trimming, aligned to the *T. oblongifolia* AmA10 reference assembly (11 Chromosomes; ~2.3 Gb) (*24*) with bwa-mem2 v2.2.1 (*25*) at default settings, and coordinate-sorted with SAMtools v1.18 (*26*). Duplicates were marked with Genome Analysis Toolkit (GATK) v4.6.1.0 MarkDuplicates (*27*). Per-sample mapping and coverage metrics were obtained with SAMtools stats and mosdepth v0.3.6 (*28*).

***Reference panel: variant calling and filtering.*** Variants were called with GATK (*29*) using HaplotypeCaller per sample, consolidated across samples with GenomicsDBImport, and jointly genotyped with GenotypeGVCFs. As no validated variant set exists for this species, variant quality score recalibration was not applied. Single-nucleotide polymorphisms (SNPs) were extracted from the joint-genotyped callset with GATK SelectVariants and hard-filtered with VariantFiltration following GATK Best Practices (QD < 2.0, FS > 60.0, SOR > 3.0, MQ < 40.0, MQRankSum < −12.5, ReadPosRankSum < −8.0) (*30*), retaining 42,660,798 of 48,313,845 raw SNP calls.

***Reference panel: population structure and differentiation.*** Filter-passing SNPs were converted to PLINK 2 v2.0.0-a.6.9LM format (*31*, *32*). One sample was excluded for genotype missingness, leaving N = 37, retaining 3,867,205 biallelic SNPs after filtering for minor allele frequency (MAF) ≥ 0.05, per-variant missingness ≤ 0.05, and per-sample missingness ≤ 0.20. Genetic differentiation among the DV, Perimeter, and Outside collection regions was estimated as Weir and Cockerham's fixation index ($F_{ST}$) (*33*). Linkage-disequilibrium (LD) decay was estimated per chromosome (Chr) as unphased $r^2$ (squared Pearson correlation coefficient) between SNP pairs within a 1 Mb window, binned by physical distance. PCA was computed from SNPs thinned by physical distance to one variant per 150 kb (13,186 SNPs). LD-based pruning was not used given the small sample size.

### Whole-genome sequencing: survival panel

***Survival panels: whole-genome sequencing.*** Libraries were prepared by the Texas A&M AgriLife Genomics and Bioinformatics Service with the PerkinElmer NEXTFLEX Rapid XP DNA-Seq Kit HT and sequenced on Illumina NovaSeq X 25B (PE150) to a target depth of ~7.5× ($N$ = 225, $n$ = 219 samples, $n$ = 6 technical replicates). Realized coverage was < 5× for 24 samples, 5–10× for 167 samples, and 10–13× for 33 samples.

***Survival panels: read processing and alignment.*** Read integrity was verified by MD5 checksum, read quality assessed with FastQC and aggregated with MultiQC, before and after trimming. Reads were concatenated across lanes per sample and trimmed with fastp with paired-end adapter auto-detection and poly-G and poly-X trimming, aligned to the *T. oblongifolia* AmA10 reference assembly (11 Chromosomes; ~2.3 Gb) (*32*) with bwa-mem2 at default settings, and coordinate-sorted with SAMtools with read groups assigned. Duplicates were marked with GATK MarkDuplicates. Per-sample mapping and coverage metrics were obtained with SAMtools stats and mosdepth.

***Survival panels: variant calling and filtering.*** Variants were called with GATK using HaplotypeCaller per sample, consolidated with GenomicsDBImport over scattered intervals, and jointly genotyped with GenotypeGVCFs; per-Chromosome VCFs were gathered with GatherVcfs. As no validated variant set exists for this species, variant quality score recalibration was not applied. SNPs were extracted from the joint-genotyped callset with GATK SelectVariants and hard-filtered with VariantFiltration following GATK Best Practices (QD < 2.0, FS > 60.0, SOR > 3.0, MQ < 40.0, MQRankSum < −12.5, ReadPosRankSum < −8.0), retaining 75,347,269 of 87,938,566 raw SNP calls.

***Survival panels: sample composition and exclusions.*** Of the $N$ = 225 libraries sequenced ($n$ = 219 individuals and $n$ = 6 technical replicates), 225 were retained in the joint-genotyped callset. Technical replicates were excluded from downstream analysis, leaving $n$ = 219 individuals assigned to collection regions as $n$ = 114 DV and Perimeter and $n$ = 105. A further nine samples were excluded for sample-level genotype missingness, giving final analysis sets of $N$ = 210 species-range, $n$ = 107 DV and Perimeter, and $n$ = 103 Outside samples.

### Imputation

***Effective population size estimate.*** Nucleotide diversity (π) was calculated in 10 kb windows with VCFtools v0.1.16 (*34*) from filter-passing SNP sets, and effective population size ($N_e$) estimated from mean π assuming a mutation rate of $7 \times 10^{-9}$ per site per generation (*35*).

***Reference panel phasing.*** Reference haplotypes were phased per Chromosome from the filter-passing SNP set of the reference panel ($N$ = 38) with BEAGLE 5.4 (*36*) in genotype-only mode, with window = 20.0, overlap = 2.0, iterations = 10 and $N_e$ = 10,000. No genetic map is available for *T. oblongifolia*, so BEAGLE's default rate of 1 cM per Mb was assumed.

***Survival panels: phasing and imputation.*** Survival-panel genotypes were phased and imputed against the phased reference panel with BEAGLE 5.4 (*36*, *37*) in two runs: a species-range run comprising all $N$ = 225 DV, Perimeter and Outside samples, and a run restricted to the $N$ = 107 DV and Perimeter samples (*38*). For each run, target genotypes were subset and restricted to sites present in the phased reference with BCFtools v1.22 (*31*), yielding 29,574,421 shared SNPs for both runs. Imputation was performed per Chromosome, with window = 40.0, overlap = 4.0, iterations = 20 with $N_e$ = 82,000 for the species-range run and $N_e$ = 85,000 for the DV/Perimeter run to retain 15,015,676 and 15,872,921 SNPs, respectively.

***Survival panels: post-imputation processing.*** Imputed variants were filtered to Dosage $R^2$ ($DR^2$) > 0.9, concatenated across Chromosomes, and restricted to biallelic SNPs with BCFtools. Dosages were converted to PLINK 2 format retaining SNPs with MAF ≥ 0.05 and per-sample missingness ≤ 0.20, with duplicate variants removed and position-sorted. The final analysis sets comprised 15,015,676 SNPs in 210 individuals for the species-range panel and 15,880,175 SNPs in 107 individuals for the DV/Perimeter panel.

***Survival panels: population structure and differentiation.*** Genetic differentiation between collection regions was estimated as $F_{ST}$ (*33*) from the imputed species-range SNP set described above. For each of species-range and DV/Perimeter SNP sets, LD decay was estimated per-chromosome as unphased $r^2$ between SNP pairs within a 2 Mb window from a random 10% subset of SNPs, binned by physical distance. As LD decayed over a shorter distance in the species-range set than in DV/Perimeter, pruning windows were set per set to retain comparable numbers of approximately independent SNPs. Species-range principal components (PCs) were computed from SNPs LD-pruned in 150 kb windows and $r^2$ > 0.2 (370,948 SNPs) (fig. S5A). DV/Perimeter PCs were computed from SNPs LD-pruned in 100 kb windows and $r^2$ > 0.2 (430,246 SNPs) (fig. S5B).

**Genome-wide association testing**

***Sample selection.*** DV/Perimeter samples passing filtering ($n$ = 107), described above, were used GWA testing with relative survival phenotypes as described below.

***Survival phenotype derivation.*** Health status scoring was as described above. A Cox proportional-hazards model (*39*) with no covariates, stratified by experimental round and tray to control for relative mortality, was fitted in the survival package v3.8.6 (*18*). Sign-inverted martingale residuals gave a continuous relative survival score that was used as the response variable in GWA testing. Positive values indicated a longer survival than expected within a stratum. Analyses were performed in R.

***Linear regression and significance testing.*** Relative survival was regressed against imputed allele dosages with PLINK 2 under an additive model, fitted per chromosome, with experimental round-by-tray indicators and PCs as covariates (Fig. 2). Models were fitted over increasing numbers of PCs, with the number retained chosen by inspection of quantile–quantile plots and the genomic inflation factor ($\lambda_{GC}$), calculated as the median observed $\chi^2$ statistic divided by its expectation under the null. PC1–25 were retained for GWA testing. Variants were checked for collinearity with covariates, with 298 of 15,880,175 SNPs excluded, leaving 15,879,877 SNPs tested. Significance was assessed against a Bonferroni threshold of α = 0.05 over the 430,246 LD-pruned variants ($P < 1.162 \times 10^{-7}$).

***Lead SNP identification.*** Passing variants were clumped in PLINK 2, associating the lowest $P$-value lead SNP with neighbors within 300 kb at squared Pearson correlation coefficient ($r^2$) > 0.1 and $P$ < 0.01 into three lead SNPs at: Chr 1 = 84,178,924 bp, Chr 2 = 178,968,191 bp, and Chr 6 = 94,545,774 bp.

***Locus boundary definition.*** For each lead SNP, unphased $r^2$ was calculated against surrounding variants in PLINK 2, and boundaries placed where LD decayed to background (Fig. 2D). Scanning outward in each direction, boundaries were extended if the preceding 100 kb contained a variant above a locus-specific $r^2$ threshold: $r^2$ = 0.4 (Chr 1), 0.175 (Chr 2) and 0.1 (Chr 6). The scan stopped once three consecutive intervals contained none. Thresholds were set per locus to reflect local LD structure: Genes within the resulting intervals were retained as candidates.

**Infrared imaging & analysis**

***Infrared data acquisition.*** IR images were acquired with an Optris PI 450 IR camera (Optris GmbH & Co) fitted with a 29° × 22° lens (f = 12.7 mm), at a detector resolution of 382 × 288 pixels and a frame rate of 27 Hz, over a measurement range of −20 to 100 °C with high-resolution temperature output enabled. Emissivity was set to 0.97 and transmissivity to 1.0. Non-uniformity correction was performed automatically at intervals of 12 to 120 s, including during recording. The camera was mounted centrally in Death Valley cabinet at 711 mm above the leaf surface, giving a field of view of ~368 × 276 mm and a pixel size of ~0.96 mm (fig. S8A). Radiometric snapshots were captured at device resolution and stored as CSV every 5 min during heat treatment and every 30 min otherwise. Because the optical properties of the IR camera lens shift as it warms, focus was pre-set before each run and left unchanged. Images were acquired only once $T_{air}$ had stabilized at the Death Valley cabinet setpoint (~30-60 min), not during the ramp.

***Leaf temperature extraction from infrared data.*** Following IR image acquisition described above, CSVs were converted to TIFF (fig. S8D). For each imaging day, one well-separated, non-overlapping leaf per plant was marked in Fiji (*41*) with the multi-point tool and saved as ROI coordinates. Leaf temperature ($T_{leaf}$) was taken as the mean of all pixels whose centers fell within a radius of 3 px (~2.9 mm) of each coordinate, applied to every snapshot from that day. Only leaves with healthy tissue were evaluated.

***Leaf trait quantification.*** Leaf outlines were traced by hand in Fiji from white-light images calibrated against a ruler in frame. Area and aspect ratio were recorded. Leaf color was expressed as green ratio (G/(R + G + B)) from mean channel intensities of the same regions. Local density was the number of adjacent tray cells containing a plant, and distance to edge the minimum number of cells to any tray edge. Traits were centered and scaled before modelling.

***Leaf trait and survival modelling.*** Multicollinearity among traits was checked by correlation and variance inflation factors. Bayesian generalized linear mixed models were fitted in MCMCglmm v2.36 (*42–45*) with tray and individual as random effects, using a categorical family for survival and a Gaussian family for leaf temperature and time to death (Fig. 3D; fig. S11, A and B). Priors were left at package defaults for fixed effects, with residual variance fixed at 1 and weakly informative priors on random effects (V = 1, v = 1). Chain length, burn-in and thinning were set per model, with chains extended until convergence was satisfied. Convergence was assessed by trace plots, autocorrelation, effective sample size, and the Heidelberger–Welch stationarity test. Analyses were performed in R.

***Leaf energy balance modelling.*** Leaf energy balance was used to model leaf-air temperature offset ($\Delta T = T_{leaf} - T_{air}$) and transpiration rate ($E$) in our extreme heat cabinet conditions by calculating net incoming and outgoing energy (*44*) (Fig. 5B; fig. S10C; fig. S11C). Constants: emissivity 0.96, Stefan–Boltzmann constant $5.67 \times 10^{-8}$ W $m^{-2}$ $K^{-4}$, latent heat of vaporization 44,000 J $mol^{-1}$, and specific heat of air 29.3 J $mol^{-1}$ $°C^{-1}$. Stomatal conductance, leaf length, and RH were each varied in turn, with remaining parameters held at values representative of the experimental heat conditions: sky temperature 55 °C, direct and diffuse shortwave radiation 275 and 20 W $m^{-2}$, atmospheric pressure 101 kPa, wind speed 0.1 m $s^{-1}$, leaf reflectance 0.65. When not the varying parameter, $T_{air}$ was constant at 60 °C, RH 0.30, stomatal conductance ($g_{sw}$) 1 mol $m^{-2}$ $s^{-1}$, and leaf length 0.017 m. Analyses were performed in R.

**Transcriptomics**

***Study design.*** Acclimation samples were taken at $T_{air}$ = 30 °C (baseline), 37 °C, 44 °C and 51 °C. At $T_{air}$ = 55.5 °C, the individuals with the lowest and highest $T_{leaf}$ were sampled separately (55_cold and 55_hot). These individuals were differentiated by IR measurement as described above during the first hour when $T_{air}$ matched Death Valley cabinet set-temperature. Developmental controls held at $T_{air}$ = 30 °C were sampled on each corresponding day (d9–d13; fig. S12A). No tissue was collected at $T_{air}$ = 60 °C, where cell-death signatures could confound the heat response.

***Leaf tissue sampling for RNA extractions.*** Seedlings were germinated and grown in standard conditions as described above. Baseline samples (30 °C) were sampled on 9 dpg, and heat acclimation was initiated on 10 dpg. One leaf from three individuals were harvested at the end of the respective 6 h heat treatment. One leaf from three developmental controls, grown in parallel in standard conditions described above, were sampled immediately after collecting heat acclimation samples. Leaves were flash frozen in liquid nitrogen followed by long-term storage at −80 °C.

***Library preparation and sequencing.*** Tissue was homogenized in 2 mL Eppendorf® tube with one 3 mm Tungsten Carbide ball in Geno/Grinder® for two rounds of 1 min at 25 strokes per second, re-frozen in between. Total RNA was extracted from with Qiagen RNeasy Plant Mini Kit (Cat no. 74904) according to manufacturer's instructions and submitted to Novogene where poly-A-selected libraries were prepared and sequenced on the Illumina NovaSeq X Plus (PE150) to a target of 6 Gb per sample (*N* = 33 libraries).

***Read processing and quantification.*** Read integrity was verified by MD5 checksum, read quality assessed with FastQC, and aggregated with MultiQC before and after trimming. Reads were trimmed with fastp with paired-end adapter auto-detection and poly-G and poly-X trimming. Trimmed reads were aligned to the *T. oblongifolia* AmA10 reference assembly (*46*) with STAR v2.7.11b (*47*), generating per-gene read counts during alignment. Counts were assembled into a gene-by-sample matrix for analysis in R.

***Sample identity verification.*** Variance-stabilized counts (DESeq2, blind) (*47*) were inspected by PCA, in which two of the six 55 °C libraries grouped with the opposite temperature condition. Each 55 °C library was then scored by the difference in Pearson correlation to the 55_hot and 55_cold centroids, excluding the library under test from both. The same two libraries scored opposite in sign to their assigned conditions; the other four were consistent. Their identities were therefore exchanged for all downstream analyses (JF11 = 55_hot and JF18 = 55_cold, initially assigned the reverse).

***Differential expression analysis.*** Transcript counts were filtered to genes with at least 10 reads in at least three samples, excluding genes on unplaced scaffolds, and differential expression (DE) tested in DESeq2 (*48*) between temperature treatments and their developmentally matched controls. Genes with adjusted $P < 0.05$ against a minimum effect size of $\log_2$ (1.5) were considered DE. $\text{Log}_2$ fold changes (L2FC) were shrunk with ashr (*49*) for multi-level contrasts and apeglm (*50*) for the direct 55 °C cold versus heat comparison. Refitting with two factors of unwanted variation estimated by the Remove Unwanted Variation (RUV) normalization strategy (*49*) gave consistent results. Analyses were performed in R.

***Developmental correction of expression.*** To separate treatment response from developmental progression, normalized counts were $\log_2$-transformed and, for each temperature, the mean expression of the treated samples was subtracted from that of the developmentally matched control (37 °C, day 10; 44 °C, day 11; 51 °C, day 12; 55 °C, day 13). The corresponding difference at the 30 °C control was subtracted as a baseline, giving a residual expression value per gene and timepoint. Analyses were performed in R.

***Gene Ontology (GO) annotation and enrichment analysis.*** GO terms were parsed from the *T. oblongifolia* annotation (*51*). Genes up-regulated in the cold- and heat-acclimated 55 °C samples were tested separately for enrichment within each GO term using clusterProfiler (*24*), against a background of all expressed genes carrying an annotation and with gene sets restricted to between 5 and 500 genes (Fig. 4C). Terms with Benjamini–Hochberg adjusted $P < 0.05$ and $q < 0.10$ were considered enriched. Analyses were performed in R.

***Gene Ontology annotation.*** Core photorespiration and auxin-binding homologs were identified by matching the Arabidopsis genes annotated with GO:0009853 (photorespiration) and GO:0010011 (auxin binding) against the ortholog assignments (*50*). *T. oblongifolia* genes were retained where the first listed Arabidopsis ortholog was one of these, and retained genes present in the expression dataset were grouped by enzyme or receptor family (fig. S12).

***Candidate gene annotation.*** Protein sequences were extracted for genes within the heat-tolerance loci (Fig. 2D) and for RNA-seq candidate genes (Fig. 4B; fig. S12). Orthologs were taken from the gene-family analysis (*52*) where available. Remaining sequences were searched against the Arabidopsis thaliana Araport11 (*53*) representative proteome with DIAMOND v2.1.8 blastp (*54*) in ultra-sensitive mode ($E < 10^{-5}$, up to 25 hits per query). Matching Arabidopsis proteins were then searched back against the *T. oblongifolia* AmA10 primary proteome ($E < 10^{-3}$, up to 5 hits per query), and genes recovering one another as best hits in both directions were designated reciprocal best hits. InterProScan v5.72-103.0 (*47*) was run on the full primary proteome with InterPro lookup and GO term retrieval enabled, and domain and GO assignments for candidate genes were taken from its output (Fig. 4, D and E; fig. S12, D and E).

***ABA application and leaf survival assay.*** Leaves were painted with 50 µM ABA in 0.05% dimethyl sulfoxide (DMSO) with 0.01% Silwet L-77, an equal volume of 0.05% DMSO with 0.01% Silwet (mock) or left untreated ~20 min before heat treatment at 60 °C. Solutions were applied to the adaxial surface with a single-use paintbrush. Each treated leaf was paired with an internal control leaf of the same developmental stage on the same plant, and both scored daily for survival as described above (Fig. 5D). Difference in time to death between treated and control leaf (Δ survival) was compared across treatments by one-way ANOVA with HSD test.

**Statistical analysis and software.**

***Use of AI-assisted technologies.*** Analysis scripts for read processing, variant calling, RNA-seq differential expression, and statistical modeling in R were written with the assistance of a large language model (Claude Sonnet 4.6, Sonnet 5, Opus 5; Anthropic). All code was reviewed, tested, and validated by the authors, and all outputs were verified against the underlying data.

***Data sources.*** WorldClim 2 (*48*), ERA5-Land (*50*) via the Open-Meteo API (*52*), Arabidopsis thaliana Araport11 release 20240409 (*53*), GBIF (*55*), iNaturalist (*21*), *T. oblongifolia* AmA10 assembly and annotation (*22*).

***Sequencing processing:*** FastQC v0.12.1 (*23*), MultiQC v1.14 (*25*), fastp v0.24.0 (*27*, *30*, *54*), bwa-mem2 v2.2.1 (*27*, *30*, *56*), GATK v4.6.1.0 (*34*, *35*), mosdepth v0.3.6 (*36*, *37*), BEAGLE 5.4 22Jul22.46e (*46*), STAR v2.7.11b (*32*).

***Sequencing analysis:*** PLINK 2 v2.0.0-a.6.9LM (*31*, *32*), BCFtools v1.22 and SAMtools v1.18 (*53*), VCFtools v0.1.16 (*54*).

***Annotation:*** DIAMOND v2.1.8 (*40*), InterProScan v5.72-103.0 (*8*).

***Imaging:*** Fiji v2.16.0 (*9*).

***R, version.*** R v4.5.2 (*57*).

***R, spatial***. rgbif v3.8.5 (*13*), rinat v0.1.10 (*14*), geodata v0.6.9 (*15*), sf v1.1.0 (*18*), terra v1.9.11 (*20*)

***R, statistics.*** lme4 v2.0.1 (*39*), lmerTest v3.2.1 (*13*), emmeans v2.0.2 (*57*), pracma v2.4.6 (*47*), survival v3.8.6 (*18*, *19*), survminer v0.5.2 (*58*), MCMCglmm v2.36 (*50*), multcompView v0.1.11 (*51*).

***R, expression.*** DESeq2 v1.50.2 (*58*), ashr v2.2.63 (*59*), apeglm v1.32.0 (*60*), RUVSeq v1.44.0 (*59*), clusterProfiler v4.18.4 (*60*).

***R, visualization.*** ggplot2 v4.0.2 (*59*), eulerr v8.1.0 (*60*), patchwork v1.3.2 (*61*), data.table v1.18.2.1 (*62*), LDheatmap v1.0.5 (*63*), ragg v1.5.2 (*64*), dplyr v1.2.0 (*65*).

14. R. Lenth, *Emmeans: Estimated Marginal Means, Aka Least-Squares Means* (2026).

15. H. Borchers, *Pracma: Practical Numerical Math Functions* (2025).

16. O. Björkman, R. W. Pearcy, A. T. Harrison, H. Mooney, Photosynthetic Adaptation to High Temperatures: A Field Study in Death Valley, California. *Science* **175**, 786–789 (1972).

17. W. H. Bennert, H. A. Mooney, The water relations of some desert plants in Death Valley, California. *Flora* **168**, 405–427 (1979).

18. T. M. Therneau, P. M. Grambsch, *Modeling Survival Data: Extending the Cox Model* (Springer, New York, 2000).

19. T. Therneau, *A Package for Survival Analysis in R* (2026).

20. A. Kassambara, M. Kosinski, P. Biecek, *Survminer: Drawing Survival Curves Using "Ggplot2"* (2026).

30. G. A. Van der Auwera, B. D. O’Connor, *Genomics in the Cloud: Using Docker, GATK, and WDL in Terra* (O’Reilly Media, Sebastopol, CA, 2020).

31. C. C. Chang, C. C. Chow, L. C. Tellier, S. Vattikuti, S. M. Purcell, J. J. Lee, Second-generation PLINK: rising to the challenge of larger and richer datasets. *Gigascience* **4**, 7 (2015).

32. S. Purcell, C. Chang, *PLINK 2.0* (www.cog-genomics.org/plink/2.0/).

43. G. S. Campbell, *An Introduction to Environmental Biophysics* (Springer, New York, NY, 1998).

44. R. W. Pearcy, J. R. Ehleringer, H. A. Mooney, P. W. Rundel, *Plant Physiological Ecology: Field Methods and Instrumentation* (Springer, Dordrecht, Netherlands, 2011).

55. iNaturalist, *iNaturalist*. https://www.inaturalist.org.

56. M. A. DePristo, E. Banks, R. Poplin, K. V. Garimella, J. R. Maguire, C. Hartl, A. A. Philippakis, G. del Angel, M. A. Rivas, M. Hanna, A. McKenna, T. J. Fennell, A. M. Kernytsky, A. Y. Sivachenko, K. Cibulskis, S. B. Gabriel, D. Altshuler, M. J. Daly, A framework for variation discovery and genotyping using next-generation DNA sequencing data. *Nat. Genet.* **43**, 491–498 (2011).

57. R Core Team, *R: A Language and Environment for Statistical Computing* (R Foundation for Statistical Computing., 2025).

58. S. Graves, H.-P. Piepho, L. Selzer, *Visualizations of Paired Comparisons [R Package multcompView Version 0.1.11]* (2026; https://CRAN.R-project.org/package=multcompView).

59. H. Wickham, *Ggplot2: Elegant Graphics for Data Analysis* (Springer International Publishing, Cham, Switzerland, ed. 2, 2016)*Use R!*

60. J. Larsson, P. Gustafsson, A Case Study in Fitting Area-Proportional Euler Diagrams with Ellipses Using eulerr. *CEUR Workshop Proceedings* **2116**, 84–91 (2018).

61. T. L. Pedersen, *Patchwork: The Composer of Plots* (2025; https://github.com/thomasp85/patchwork).

62. T. Barrett, M. Dowle, A. Srinivasan, J. Gorecki, M. Chirico, T. Hocking, B. Schwendinger, I. Krylov, *Data.Table: Extension of `data.Frame`* (2026; https://CRAN.R-project.org/package=data.table).

63. J.-H. Shin, S. Blay, J. Graham, B. McNeney, LDheatmap: An R Function for Graphical Display of Pairwise Linkage Disequilibria Between Single Nucleotide Polymorphisms. *J. Stat. Softw.* **16**, 1–9 (2006).

64. T. L. Pedersen, M. Shemanarev, *Ragg: Graphic Devices Based on AGG* (2026).

65. H. Wickham, R. François, L. Henry, K. Müller, D. Vaughan, *Dplyr: A Grammar of Data Manipulation* (2026; https://dplyr.tidyverse.org).
