## supplementary figures and tables for "Extreme cooling enables survival in extreme heat"

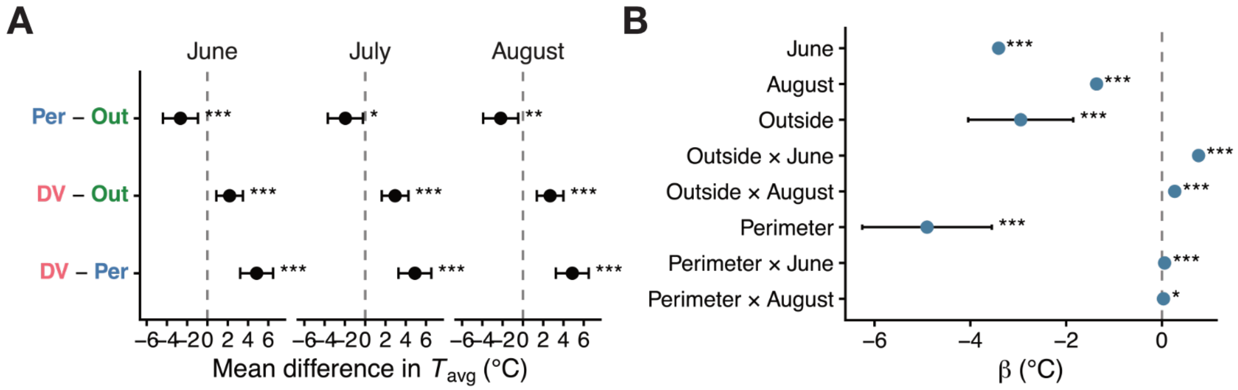


**fig. S1 Death Valley climates are significantly hotter than both Perimeter and Outside climates.** Statistical model underlying Fig. 1C. Summer daytime average air temperature ($T_{avg}$; 1100–2100) at seed collection site coordinates by collection region: Death Valley (DV), Perimeter (Per), Outside (Out) and month. **(A)** Pairwise differences in mean $T_{avg}$ between regions, estimated from a linear mixed model [lmer($T_{avg}$ ~ region × month + hour + (1 | site))]. Contrasts were computed as estimated marginal means with Tukey adjustment; points are mean difference (°C), bars 95% CI. Positive values indicate the first region is warmer. **(B)** Fixed-effect coefficients (β, °C) from the model in (A), relative to DV in July; bars are 95% CI. The effect of region was significant [$F_{(2,37)} = 28.1$, $P = 3.76 \times 10^{-8}$]. Asterisks: *$P < 0.05$, **$P < 0.01$, ***$P < 0.001$.

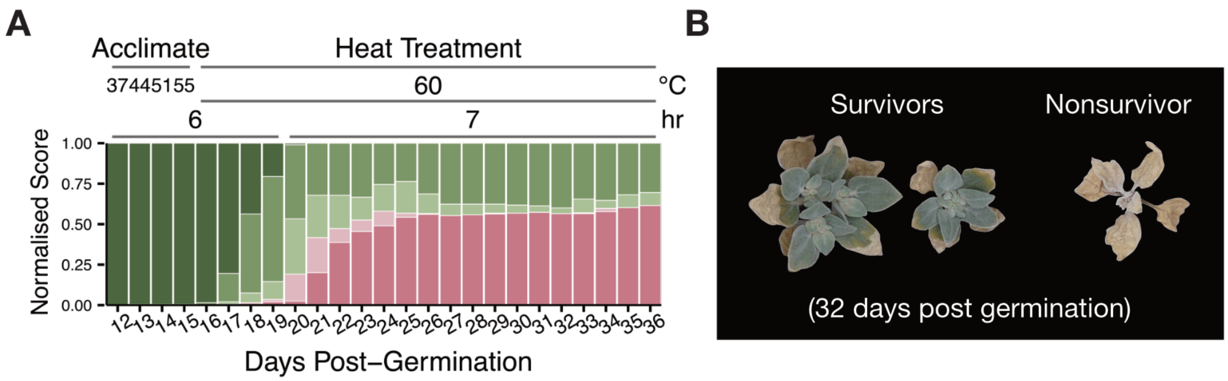


**fig. S2 Some genotypes of *T. oblongifolia* can survive at least 20 days at 60 °C. (A)** Distribution of daily health status scores across extreme heat survival assay for Fig. 3G. Plants were acclimated in stepwise 6 h increments (37, 44, 51, 55 °C) then held at 60 °C for 7 h daily. Bars show the proportion of plants in each score category (green = healthy, pink = dead) per day post-germination (dpg) (methods). **(B)** Representative survivors and a nonsurvivor at 36 dpg**.**

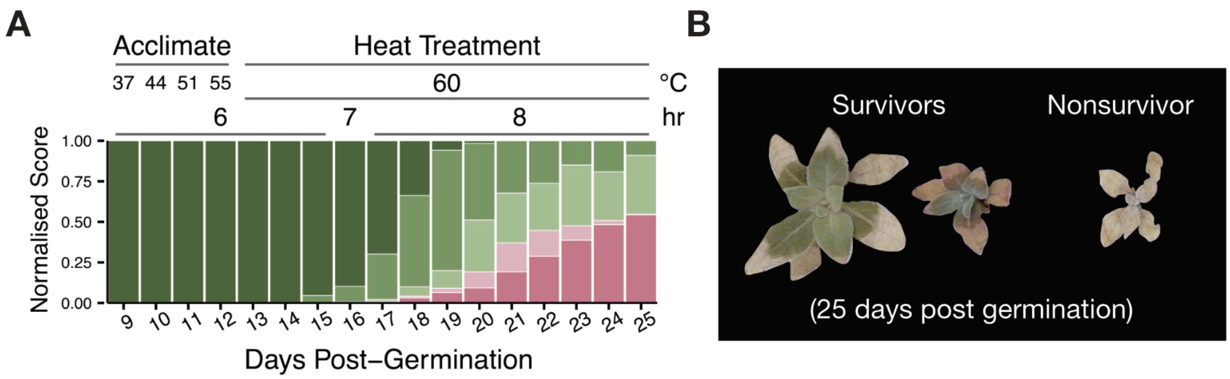


**fig. S3 Progeny from wild *T. oblongifolia* seed grown in greenhouse survive at 60 °C. (A)** Distribution of daily health status scores across extreme heat survival assay. Plants were acclimated in stepwise 6 h increments (37, 44, 51, 55 °C), held at 60 °C for 7 h on the first day and 8 h daily thereafter. Bars show the proportion of plants in each score category (green = healthy, pink = dead) per dpg (methods). **(B)** Representative survivors and a nonsurvivor at 25 dpg. $n$ = 313.

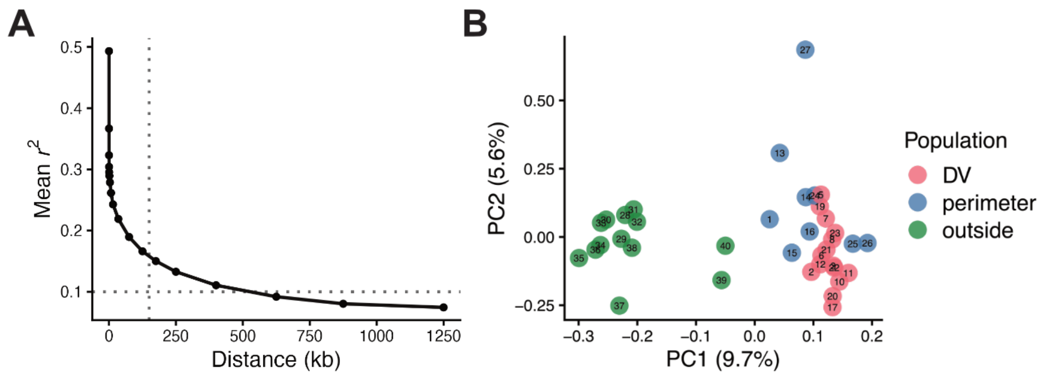


**fig. S4 Population substructure of *T. oblongifolia* reference panel across species range. (A)** Linkage disequilibrium (LD) defined by $r^2$ (squared Pearson correlation coefficient) decays rapidly. Mean LD against physical distance between SNP pairs, binned and averaged across chromosomes in PLINK 2 (--r2-unphased). Dotted lines mark $r^2$ = 0.1 (horizontal) and 150 kb (vertical). **(B)** PCA of distance-thinned (see methods), hard-filtered single-nucleotide polymorphisms (SNPs) across all individuals, colored by collection region. 13,186 SNPs retained after thinning to one variant per 150 kb. Points are individuals; numbers label site ID centroids. Weir and Cockerham estimation of fixation index ($F_{ST}$) = 0.0563 (Outside and DV), 0.0487 (Outside and Perimeter), 0.00611 (DV and Perimeter).

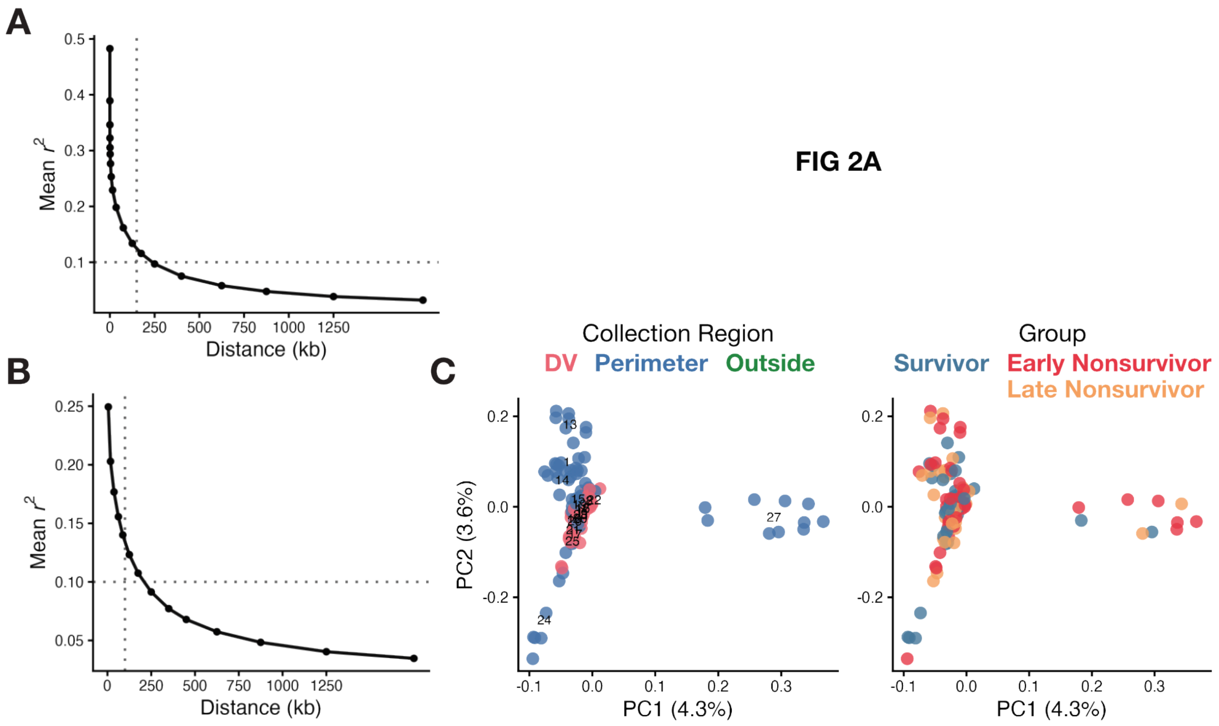


**fig. S5 Population substructures from whole-genome sequencing of extreme heat survivor samples.** **(A)** LD pruning window ahead of the PCA in Fig. 2A. LD defined by $r^2$ decays rapidly for imputed SNPs across the full imputed survival panel. Mean LD against physical distance between SNP pairs, binned and averaged across chromosomes in PLINK 2 (--r2-unphased). Dotted lines mark $r^2$ = 0.1 (horizontal) and 150 kb (vertical). **(B)** LD pruning window ahead of PCA in (C). LD defined by $r^2$ decays rapidly as in (A). Dotted lines mark $r^2$ = 0.1 (horizontal) and 100 kb (vertical). **(C)** DV/Perimeter are not differentiated by population substructure nor survival phenotypes. PCA of the independently imputed DV/Perimeter SNPs, LD-pruned in 100 kb windows at $r^2 < 0.2$ (430,246 SNPs retained), colored by collection region (left) and survival group (right). Points are individuals; numbers label site ID centroids.

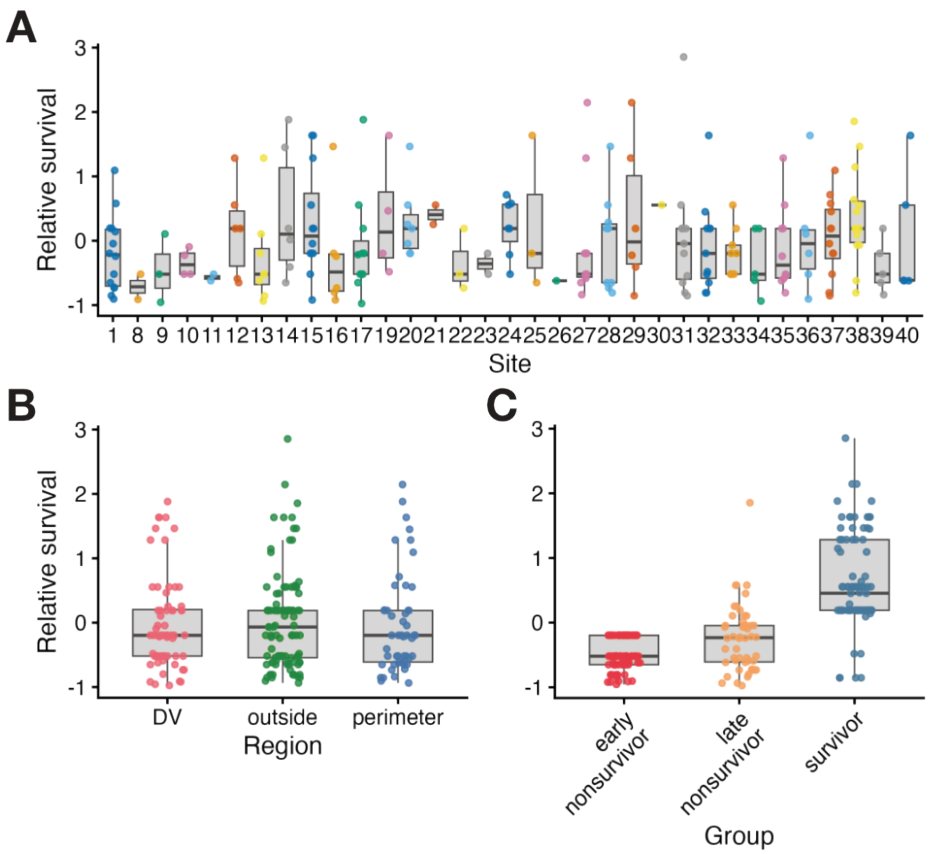


**fig. S6 Relative survival is not structured by geography.** Relative survival (methods) of whole-genome sequenced extreme heat survival-assay individuals (n = 219; table S3), by **(A)** collection site, **(B)** collection region (DV, Perimeter, Outside), and **(C)** survival group, assigned from *k*-means clustering of daily health-score trajectories (methods). Points are individuals; boxes, median and interquartile range (IQR); whiskers, 1.5 × IQR. Relative survival differed among survival groups (Kruskal–Wallis $P = 1.15 \times 10^{-25}$) but not among collection sites ($P = 0.365$) or regions ($P = 0.718$).

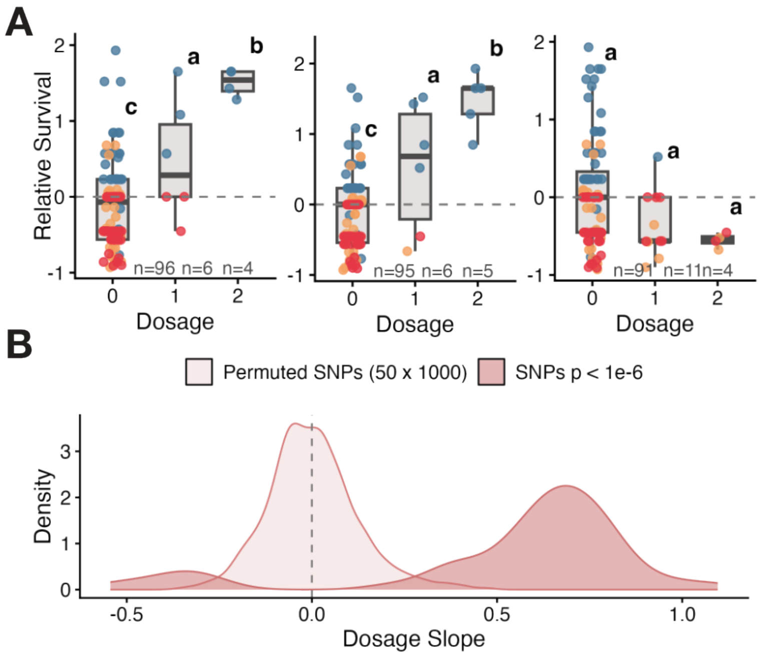


**fig. S7 Alternate allele dosage at associated loci corresponds to survival. (A)** Relative survival by alternate allele dosage at three lead SNPs. Points are individuals colored by survival group; boxes, median and IQR; whiskers, 1.5 × IQR. Letters denote dosage classes differing significantly (ANOVA with Tukey HSD; Chr1 $P = 3.75 \times 10^{-7}$, Chr2 $P = 7.20 \times 10^{-9}$, Chr 6 $P = 0.040$). Sample sizes per class are given below each box. **(B)** Distribution of per-SNP dosage slopes (regression of relative survival on dosage) for SNPs associated with survival at $P < 1.0 \times 10^{-6}$, compared with a null distribution of means from 1,000 random draws of 50 SNPs sampled genome-wide. Dashed line, slope = 0.

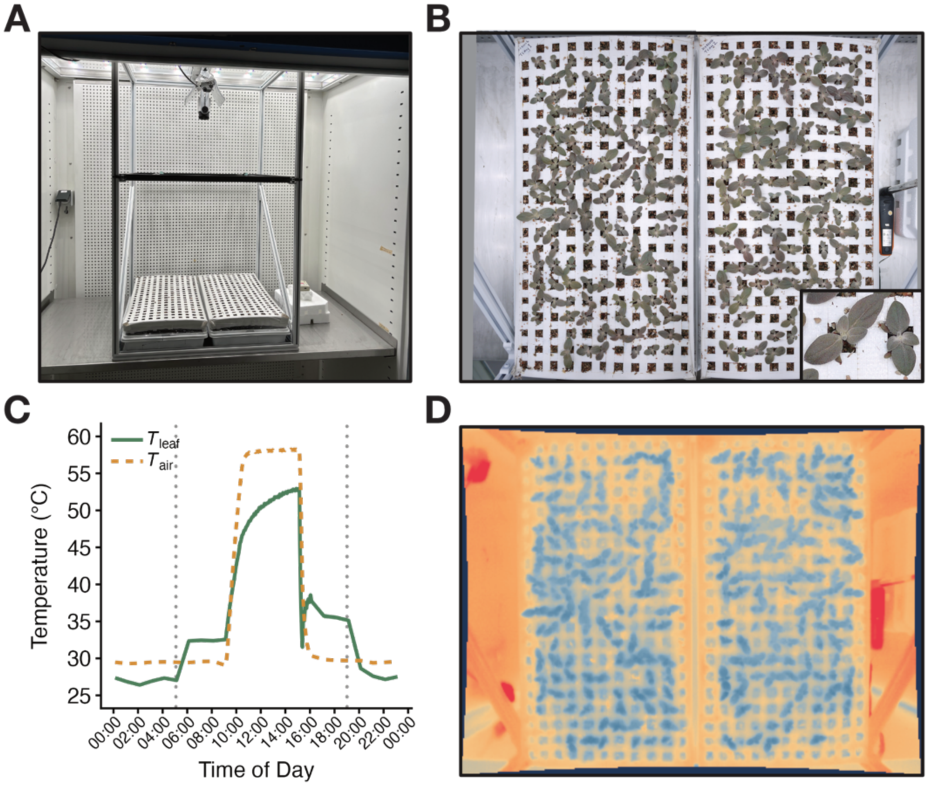


**fig. S8 High-throughput Infrared imaging of *T. oblongifolia* seedlings at extreme temperatures.** **(A)** Growth cabinet with plant trays and overhead Infrared (IR) camera. **(B)** Overhead visible-light image of two trays of *T. oblongifolia seedlings*. Inset is close-up white-light image of *T. oblongifolia* seedlings. **(C)** $T_{air}$ and a representative mean leaf temperature ($T_{leaf}$) over a 24 h cycle. Dotted grey lines, photoperiod is 14 h from 0500-1900. **(D)** Representative overhead IR image of the same trays as in (B), showing leaves cooler than the surrounding tray surface.

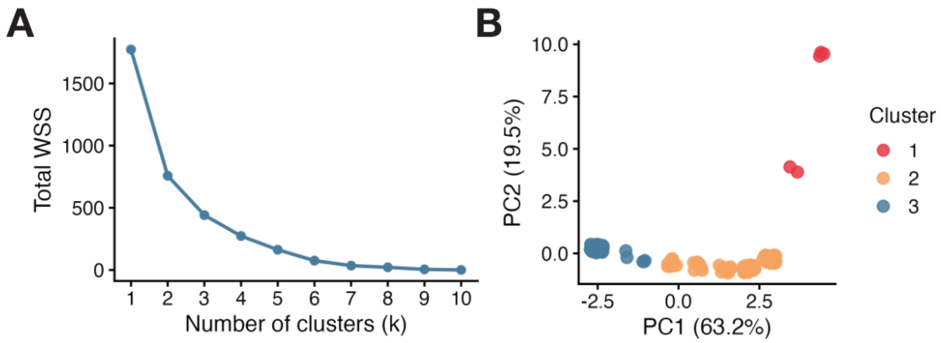


**fig. S9 Clustering of health-score trajectories. (A)** Total within-cluster sum of squares against number of clusters (*k*) for *k*-means clustering of normalized daily health scores across 60 °C heat treatment. **(B)** PCA of the same normalized scores, colored by *k*-means cluster (*k* = 3). Data for Fig. 3, A to D.

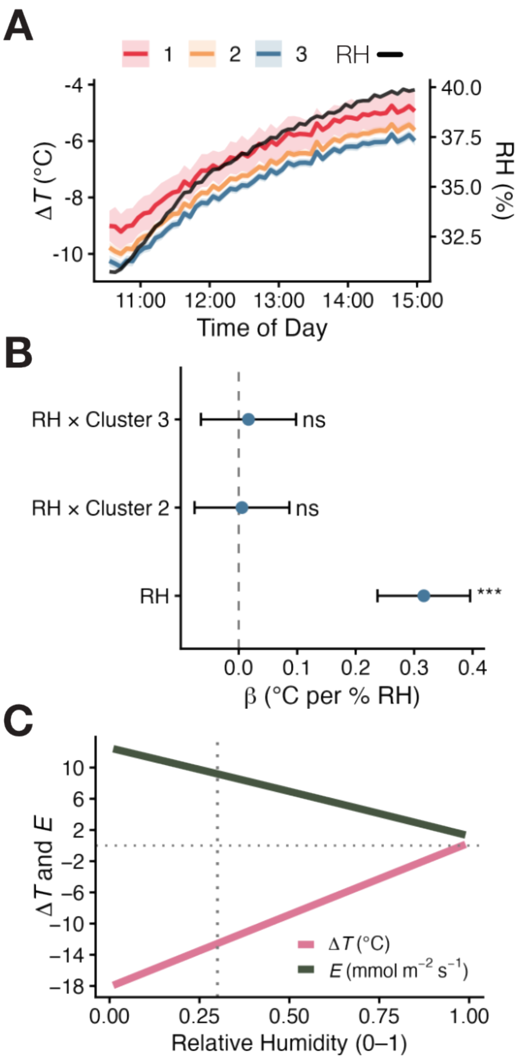


**fig. S10 Rising relative humidity correlates with rise in less negative leaf cooling. (A)** Data as in Fig. 3B with relative humidity (RH) overlaid (RH, black; right axis). **(B)** Fixed-effect coefficients (β, °C per % RH) from a linear mixed model, lmer($\Delta T$ ~ RH × cluster + (1 | cell)); points, estimates; bars, 95% CI. Cluster 1 is the model reference level. RH affected $\Delta T$ but did not differ between clusters. ****P* < 0.001; ns, not significant. **(C)** Modelled $\Delta T$ and transpiration rate (*E*) against RH, from energy balance modelling (see materials and methods). Dotted lines, cabinet setpoint (RH = 0.30) and $\Delta T$ = 0.

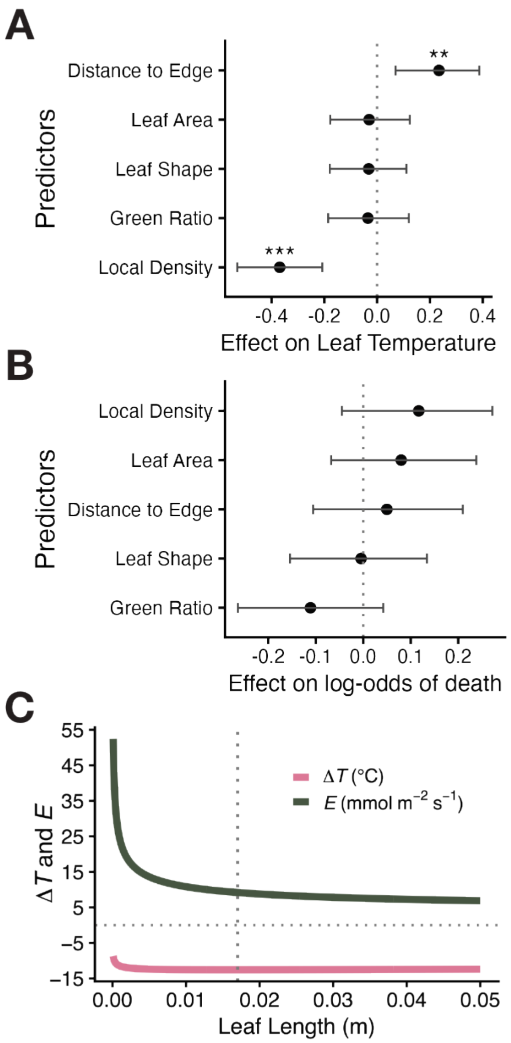


**fig. S11 Leaf position, not morphology, predicts leaf temperature, and no trait predicts death independently of temperature. (A)** Effect of leaf traits on standardized $T_{leaf}$, estimated by Bayesian generalized linear mixed model (MCMCglmm, Gaussian) with tray and individual as random effects. Predictors and response were standardized; estimates are per 1 SD. Points, posterior means; bars, 95% credible intervals. Local density and distance to edge had credible effects (**pMCMC < 0.01, ***pMCMC < 0.001). **(B)** As in (A), for effect on log-odds of death (MCMCglmm, categorical), fitted without leaf temperature as a predictor. No trait had a credible effect. **(C)** Modelled $\Delta T$ and transpiration rate ($E$) against leaf length, from leaf energy balance. Dotted vertical line, mean *T. oblongifolia* leaf length (0.017 m); dotted horizontal line, $\Delta T = 0$.

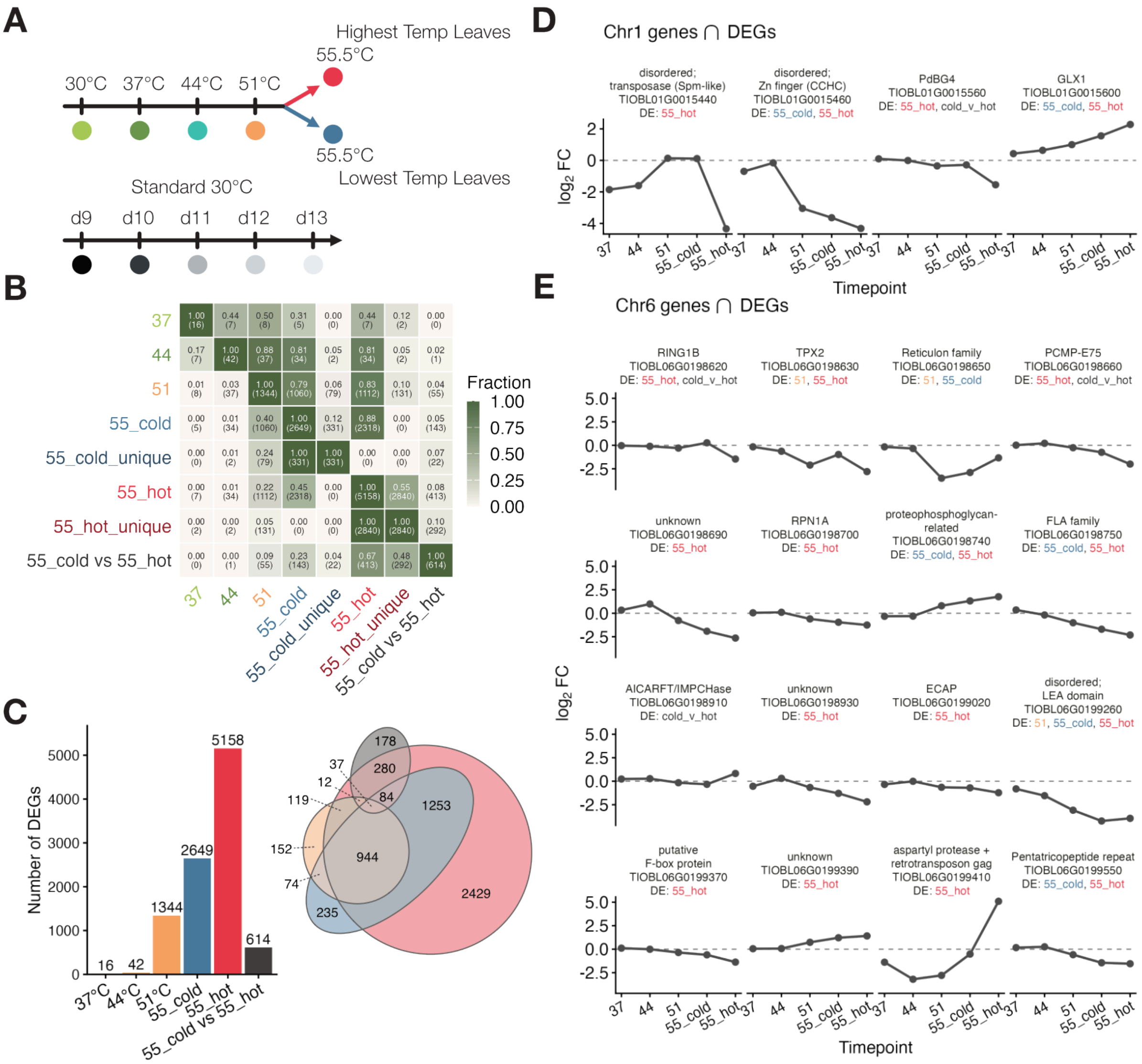


**fig. S12 Transcriptional response to heat treatment across timepoints and GWAS candidate genes.** **(A)** Sampling scheme for RNAseq. Plants were sampled at 30, 37, 44, and 51 °C, then at 55.5 °C split into the highest (55_hot) and lowest (55_cold) leaf-temperature samples. Day-matched controls held at 30 °C were sampled at 9–13 days post-germination (d9–d13). **(B)** Pairwise overlap between differentially expressed gene (DEG) sets. Color and value give the fraction of the row set contained in the column set; counts in parentheses. Sets are DEGs at each temperature relative to day-matched controls (37, 44, 51, 55_cold, 55_hot), genes significant in one 55 °C group but not the other (55_cold_unique, 55_hot_unique), and genes differentially expressed between 55_cold vs 55_hot. **(C)** Left: number of DEGs per contrast. Right: Euler diagram of overlap among 51 °C, 55_cold, 55_hot, 55_cold vs 55_hot. **(D)** $\log_2$ fold change (L2FC) across timepoints for genes at Chr 1 that are also DEGs. Gene annotation, identifier, and the contrasts in which each is differentially expressed are shown above each panel. **(E)** As in (D), for Chr 6. Combined, 19 candidate genes at heat tolerance loci are present in 51, 55_cold and 55_hot DEGs, with four in 55_cold versus 55_hot DEGs.

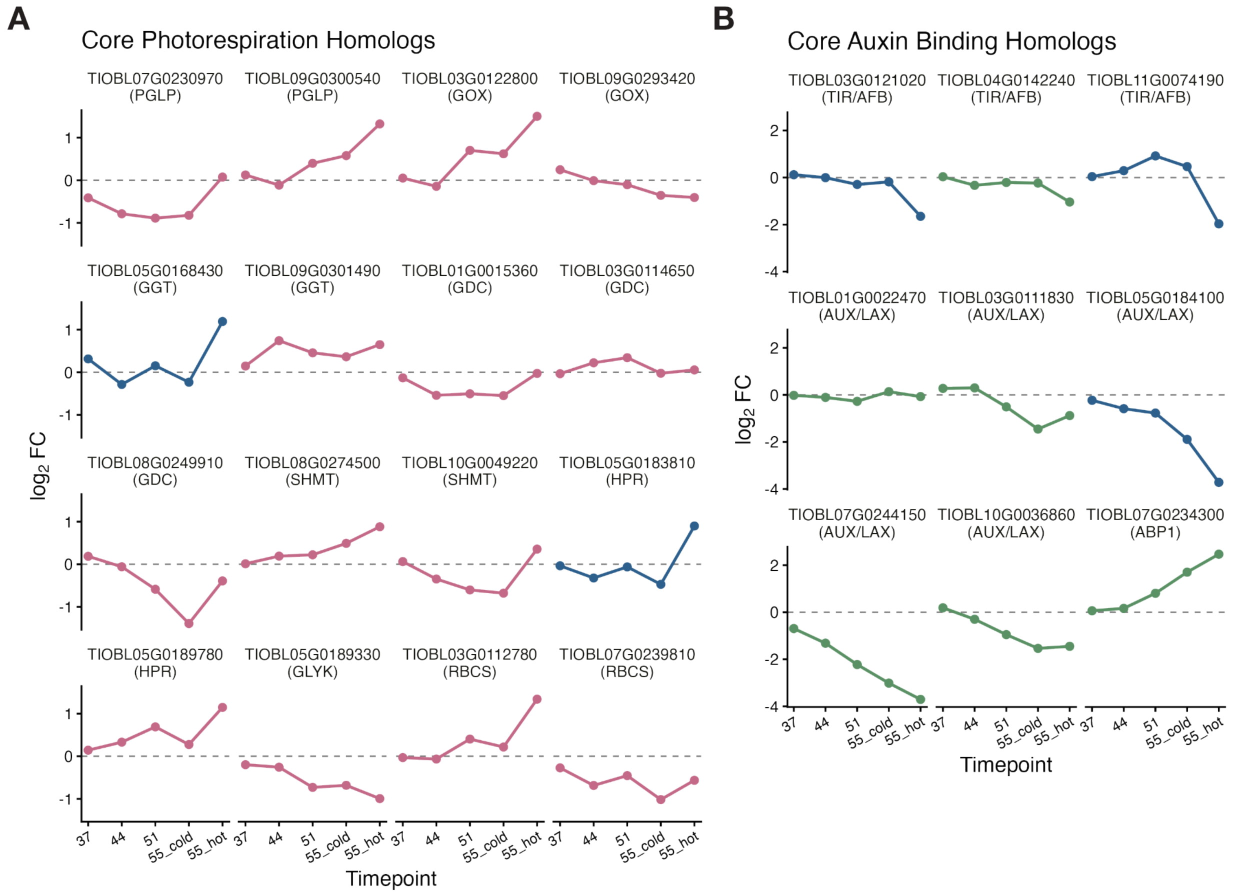


**fig. S13 Expression of core photorespiration and auxin binding homologs across heat treatment. (A)** L2FC change relative to day-matched controls across timepoints for *T. oblongifolia* homologs of core Arabidopsis photorespiration genes, grouped by gene family. Blue, homologs among the genes driving enrichment of that GO term (Fig. 4C); pink, additional pathway homologs not recovered by the enrichment. Dashed line, L2FC = 0. **(B)** As in (A), for core auxin binding homologs. Blue, genes in the auxin binding GO term enriched among genes up in 55_cold (Fig. 4C); green, additional pathway homologs not recovered by the enrichment.

| Collection Trip | Site ID | Site Description | Collection Region | Coordinates (DD, WGS84) | Individuals |
|---|---|---|---|---|---|
| 202302 DEVA | 1 | spring meadows | perimeter | 36.42548, -116.38719 | 11 |
| 202302 DEVA | 2 | zabrinski point | DV | 36.41621, -116.80384 | 6 |
| 202302 DEVA | 3 | furnace south | DV | 36.44303, -116.85135 | 6 |
| 202302 DEVA | 4 | badwater north | DV | 36.40314, -116.84891 | 1 |
| 202302 DEVA | 5 | badwater north | DV | 36.38607, -116.85214 | 6 |
| 202302 DEVA | 6 | artist | DV | 36.33060, -116.82766 | 6 |
| 202302 DEVA | 7 | artist | DV | 36.34976, -116.79326 | 6 |
| 202302 DEVA | 8 | badwater north | DV | 36.38424, -116.83897 | 6 |
| 202302 DEVA | 9 | badwater south | DV | 36.26930, -116.78637 | 6 |
| 202302 DEVA | 10 | badwater south | DV | 36.22474, -116.77755 | 6 |
| 202302 DEVA | 11 | badwater south | DV | 36.16080, -116.76714 | 6 |
| 202302 DEVA | 12 | badwater south | DV | 36.03929, -116.75909 | 6 |
| 202302 DEVA | 13 | hw127 | perimeter | 36.16753, -116.32489 | 4 |
| 202302 DEVA | 14 | spring meadows | perimeter | 36.40079, -116.42247 | 6 |
| 202302 DEVA | 15 | hw190 east | perimeter | 36.30388, -116.45941 | 6 |
| 202302 DEVA | 16 | hw190 west | perimeter | 36.37459, -116.69403 | 6 |
| 202302 DEVA | 17 | furnace south | DV | 36.44869, -116.85531 | 6 |
| 202302 DEVA | 18 | the ranch | DV | 36.45831, -116.86558 | 1 |
| 202302 DEVA | 19 | furnace north | DV | 36.47444, -116.86763 | 6 |
| 202302 DEVA | 20 | furnace north | DV | 36.51072, -116.88048 | 6 |
| 202302 DEVA | 21 | stovepipe | DV | 36.58962, -116.94869 | 6 |
| 202302 DEVA | 22 | stovepipe | DV | 36.62252, -117.06096 | 6 |
| 202302 DEVA | 23 | stovepipe | DV | 36.58309, -117.17559 | 5 |
| 202302 DEVA | 24 | panamint | perimeter | 36.25747, -117.22301 | 6 |
| 202302 DEVA | 25 | panamint | perimeter | 36.33916, -117.47984 | 6 |
| 202302 DEVA | 26 | panamint | perimeter | 36.34283, -117.37665 | 5 |
| 202302 DEVA | 27 | beatty | perimeter | 36.86548, -116.75608 | 6 |
| 202303 SOCAL | 28 | hw8 | outside | 32.74833, -115.92547 | 6 |
| 202303 SOCAL | 29 | anzaborrego | outside | 32.85689, -116.20668 | 4 |
| 202303 SOCAL | 30 | borregosprings | outside | 33.30035, -116.26866 | 2 |
| 202303 SOCAL | 31 | borregosprings | outside | 33.28193, -116.14111 | 6 |
| 202303 SOCAL | 32 | salton city | outside | 33.17042, -115.87972 | 6 |
| 202303 SOCAL | 33 | blythe | outside | 33.30671, -114.74207 | 6 |
| 202303 SOCAL | 34 | quartz | outside | 33.69743, -114.21694 | 6 |
| 202303 SOCAL | 35 | parker | outside | 33.99582, -114.22043 | 6 |
| 202303 SOCAL | 36 | havasu | outside | 34.29023, -114.10368 | 5 |
| 202303 SOCAL | 37 | pht | outside | 34.72043, -114.45187 | 6 |
| 202303 SOCAL | 38 | bullhead city | outside | 35.11061, -114.54494 | 6 |
| 202303 SOCAL | 39 | lake mead | outside | 35.97886, -114.69775 | 6 |
| 202303 SOCAL | 40 | lake mead | outside | 36.11117, -114.86001 | 5 |

**table S1** *Tidestromia oblongifolia* field collection sites. Seed was collected from 40 sites across the species range during two trips in 2023: Death Valley National Park and surrounding areas in late February (202302 DEVA, sites 1–27) and southern California in early March (202303 SOCAL, sites 28–40). Sites were assigned to one of three collection regions: Death Valley (DV), Perimeter, or Outside. Individuals gives the number of maternal plants sampled per site (223 total).

| Region | Seedling | Site | Individual | Individual/ Site | Seedling/ Total | Site/ Total | Individual/ Total |
|---|---|---|---|---|---|---|---|
| DV | 246 | 14/16 | 31/94 | 0.308 | 0.194 | 0.389 | 0.244 |
| Perimeter | 390 | 9/9 | 39/56 | 0.703 | 0.308 | 0.250 | 0.307 |
| Outside | 632 | 13/13 | 57/70 | 0.820 | 0.498 | 0.361 | 0.449 |
| Total | 1268 | 36/38 | 127 | - | - | - | - |

**table S2** Screening counts by collection region. Seedling, seedlings screened. Site and Individual, number screened out of the number collected. Individual/Site, proportion of collected individuals that were screened. Seedling/Total, Site/Total, and Individual/Total, each region's share of the corresponding total.

| Group | DV | Perimeter | Outside | Total |
|---|---|---|---|---|
| Survivor | 22 | 15 | 50 | 87 |
| Early Nonsurvivor | 28 | 20 | 34 | 82 |
| Late Nonsurvivor | 10 | 19 | 21 | 50 |
| Total | 60 | 54 | 105 | 219 |

**table S3** Survival outcomes by collection region. Plants were classified as survivors, early nonsurvivors, or late nonsurvivors from *k*-means clustering of daily health-score trajectories.

| Region 1 | Region 2 | $F_{ST}$ |
| --- | --- | --- |
| Outside | DV | 0.0616 |
| Outside | Perimeter | 0.0621 |
| DV | Perimeter | 0.0193 |

**table S4** $F_{ST}$ between collection regions among whole-genome sequenced survival assay individuals (Fig. 2), calculated from imputed SNPs (15,015,676).
